# Effectors enabling adaptation to mitochondrial complex I loss in Hürthle cell carcinoma

**DOI:** 10.1101/2022.08.16.504041

**Authors:** Raj K. Gopal, Venkata R. Vantaku, Apekshya Panda, Bryn Reimer, Sneha Rath, Tsz-Leung To, Adam S. Fisch, Murat Cetinbas, Maia Livneh, Michael J. Calcaterra, Benjamin J. Gigliotti, Kerry Pierce, Clary B. Clish, Dora Dias-Santagata, Peter M. Sadow, Lori J. Wirth, Gilbert H. Daniels, Ruslan I. Sadreyev, Sarah E. Calvo, Sareh Parangi, Vamsi K. Mootha

## Abstract

Oncocytic (Hürthle cell) carcinoma of the thyroid (HCC) is genetically characterized by complex I mitochondrial DNA mutations and widespread chromosomal losses. Here, we utilize RNA-seq and metabolomics to identify candidate molecular effectors activated by these genetic drivers. We find glutathione biosynthesis, amino acid metabolism, mitochondrial unfolded protein response, and lipid peroxide scavenging, a safeguard against ferroptosis, to be increased in HCC. A CRISPR-Cas9 knockout screen in a new HCC model reveals which pathways are key for fitness, and highlights loss of GPX4, a defense against ferroptosis, as a strong liability. Rescuing complex I redox activity with the yeast NADH dehydrogenase (NDI1) in HCC cells diminishes ferroptosis sensitivity, while inhibiting complex I in normal thyroid cells augments ferroptosis induction. Our work demonstrates unmitigated lipid peroxide stress to be an HCC vulnerability that is mechanistically coupled to the genetic loss of mitochondrial complex I activity.

**Significance:** Oncocytic (Hürthle cell) carcinoma of the thyroid (HCC) is a unique tumor with a remarkable accumulation of mitochondria. HCC harbors unique genetic alterations, including mitochondrial DNA (mtDNA) mutations in complex I genes and widespread loss-of-heterozygosity in the nuclear DNA. With less favorable clinical outcomes, new therapies for HCC are needed, especially since these tumors show intrinsic resistance to radioactive iodine, which is one of the main treatments for metastatic well-differentiated thyroid cancer. An absence of authentic HCC cell lines and animal models has hindered the mechanistic understanding of this disease and slowed therapeutic progress. In this study, we describe the transcriptomic and metabolomic landscape of HCC and present new HCC models that recapitulate key mtDNA and nuclear DNA alterations. A targeted CRISPR-Cas9 knockout screen in an HCC cell line highlights the molecular programs nominated by our -omics profiling that are required for cell fitness. This screen suggests that lipid peroxide scavenging, a defense system against an iron-dependent form of cell death known as ferroptosis, is a vulnerability in HCC that is coupled to complex I loss, and that targeting this pathway may help patients with HCC.

## Introduction

Oncocytic (Hürthle cell) carcinoma of the thyroid (HCC) has a tendency for aggressive behavior and an intrinsic resistance to radioactive iodine (RAI), a central therapy for patients with well-differentiated thyroid cancer (Besic et al., 2003). HCC is a unique subtype of differentiated thyroid cancer in that it has a notable abundance of dysmorphic mitochondria, a histological morphology seen in oncocytic neoplasms (Maximo and Sobrinho-Simoes, 2000). While oncocytic tumors of other anatomic sites, such as renal oncocytoma (RO), are predominantly indolent, HCC can have distant metastases with strong uptake of fluorodeoxyglucose on PET imaging (Lowe et al., 2003).

The genetic underpinnings of HCC include complex I mitochondrial DNA (mtDNA) mutations and widespread chromosomal losses. While early targeted studies identified mtDNA mutations and marked aneuploidy as characteristic of HCC (Corver et al., 2012; Gasparre et al., 2007; Maximo et al., 2002; Mazzucchelli et al., 2000; Tallini et al., 1999), the role of these events was difficult to contextualize in the absence of a comprehensive genetic analysis. Two recent whole exome HCC studies established that mtDNA mutations and widespread chromosomal losses are the defining driver events in HCC(Ganly et al., 2018; Gopal et al., 2018b). The mtDNA mutations are disruptive and significantly enriched in complex I genes, suggesting loss of the first step in mitochondrial oxidative phosphorylation (OXPHOS) similar to other oncocytic tumors (Gopal et al., 2018a; Mayr et al., 2008; Simonnet et al., 2003). In the nuclear genome, chromosomal loss events result in extensive loss-of-heterozygosity (LOH), culminating in either a near-haploid or uniparental disomic state. A detailed phylogenetic analysis revealed that widespread LOH and mtDNA complex I mutations are truncal events maintained during tumor evolution, further supporting their role in tumor initiation and progression (Gopal et al., 2018b).

The core molecular effectors activated in the face of the mitochondrial and nuclear genomic alterations of HCC have not been well characterized. Efforts to uncover these molecular pathways and move towards implementing novel therapeutic strategies have stalled due to an absence of reliable HCC models. Existing gene expression and metabolic analyses of HCC have identified changes in the tricarboxylic acid cycle as well as activation of the mTOR pathway (Addie et al., 2020; Dong et al., 2022; Ganly et al., 2022; Ganly et al., 2013). The growth-promoting and adaptive pathways activated by loss of mitochondrial complex I, however, have not been well characterized in HCC.

To address these questions, we perform joint RNA-seq and metabolomic profiling of an HCC cohort with matched normal thyroid samples and find coordinated upregulation in amino acid and glutathione biosynthesis as well as the accumulation of lipids with polyunsaturated fatty acids (PUFAs) in tumors. A CRISPR-Cas9 screen together with targeted drug screening validates the essentiality of these metabolic pathways in newly created authentic HCC disease models. We further identify lipid peroxide stress as a novel vulnerability that is mechanistically coupled to mitochondrial complex I loss and that can potentially be exploited to treat HCC patients.

## Results

### An HCC cohort with complex I mtDNA mutations and chromosomal losses

With the goal of performing RNA-seq and metabolomic profiling, we established a new cohort of fresh frozen HCC samples and confirmed their mtDNA mutation status and chromosomal copy number. This cohort, which was almost entirely distinct from our recent whole exome analysis (Gopal et al., 2018b), contained 24 oncocytic (Hürthle cell) tumors with 18 cases having matched normal thyroid tissue (Supplementary Table 1). Tumor samples included 21 primary tumors comprised of 19 HCC (8 widely invasive, 11 minimally invasive) and 2 oncocytic (Hürthle cell) adenomas as well as 2 locoregional recurrences (LR) and 1 distant metastasis (DM; Figure 1A).

**Figure 1:**
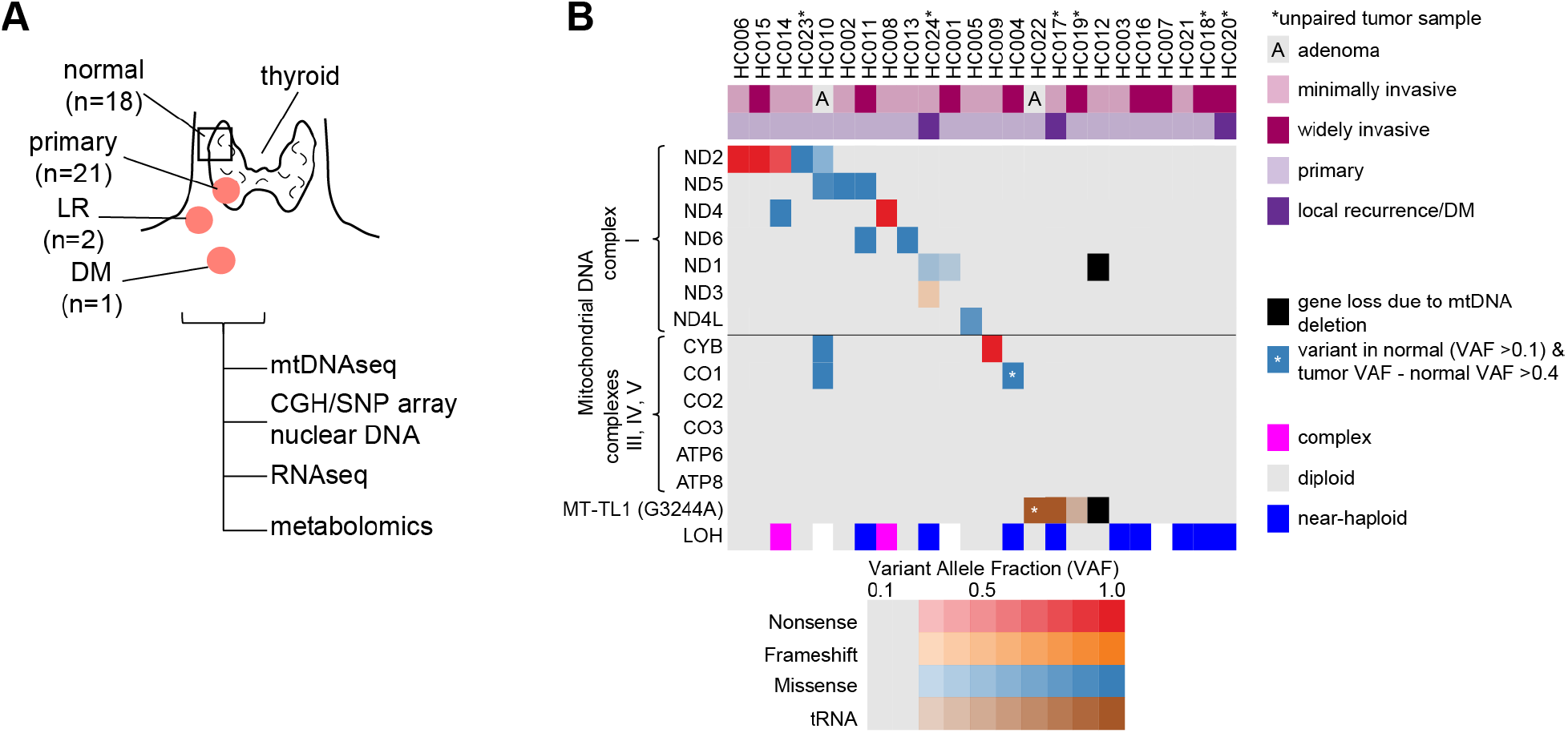
An HCC cohort with complex I mtDNA mutations and chromosomal losses. (A) Clinical cohort of HCC and normal thyroid samples with profiling methods used. Primary, primary tumor; LR, locoregional recurrence; DM, distant metastasis. (B) Mutations in mtDNA organized by complex with allelic fraction > 0.2 with CGH-SNP array copy number profile. LOH, loss-of-heterozygosity. See also Figure S1.

Next-generation mtDNA sequencing identified 18 of the 24 tumors (75%) to harbor a mtDNA alteration at >0.20 variant allele fraction (VAF), a surrogate of heteroplasmy (Figure 1B; Supplementary Table 2). There were 14 tumors with disruptive mtDNA mutations in OXPHOS encoding genes, including 12 complex I, one complex III, and one complex IV mutant tumors (Figure 1B). Three tumors contained a tRNA variant in *MT-TL1* (G3244A) previously associated (Kobayashi et al., 1987) with mitochondrial encephalomyopathy, lactic acidosis, and stroke-like episodes (MELAS), a mitochondrial disorder associated with complex I deficiency (Figure 1B). Finally, one tumor contained a 3710 bp deletion that led to loss of a complex I gene (*MT-ND1*) and *MT-TL1* (Figure 1B and Supplementary Figure 1A). We confirmed these mtDNA alterations in 16 of 18 samples using RNA-seq and found concordant variant calls in three patients that were part of our prior exome cohort (Supplementary Table 2).

We used array-based comparative genomic hybridization CGH (aCGH) combined with SNP microarrays to determine the nuclear DNA copy number state in our sample cohort. The chromosomal state was identified in 21 of 24 samples with the remaining 3 samples not passing quality control. Tumors existed in 3 copy number profiles: “diploid” (n=10), “near-haploid” (n=9), and “complex” (n=2; Figure 1B, Supplementary Figure 1B, and Supplementary Table 2). All 10 diploid tumors harbored a disruptive OXPHOS alteration. In addition, of the 6 tumors with no mtDNA changes, the 5 with reliable aCGH and SNP array data were all near-haploid (Figure 1B).

Overall, our new cohort of HCC patients, which we established for RNA-seq and metabolomic profiling, recapitulates the key nuclear DNA and mtDNA findings that are predicted based on recent exome analyses of HCC (Ganly et al., 2018; Gopal et al., 2018b).

### Transcriptional landscape of HCC

We performed bulk RNA-seq on our entire cohort to identify differentially expressed molecular programs in HCC. Principal component analysis (PCA) showed all tumors, regardless of mtDNA or chromosomal status, clustered along the first principal component (PC1; 22.5% of the variance; Figure 2A). The 4 samples diverging along the second principal component (PC2) had Hashimoto thyroiditis with PC2 loadings comprised of immunoglobulin genes (Figure 2A; Supplementary Table 3). We observed significant differences in gene expression with 1987 upregulated and 4337 downregulated transcripts (adjusted p < 0.001; Supplementary Table 3).

**Figure 2:**
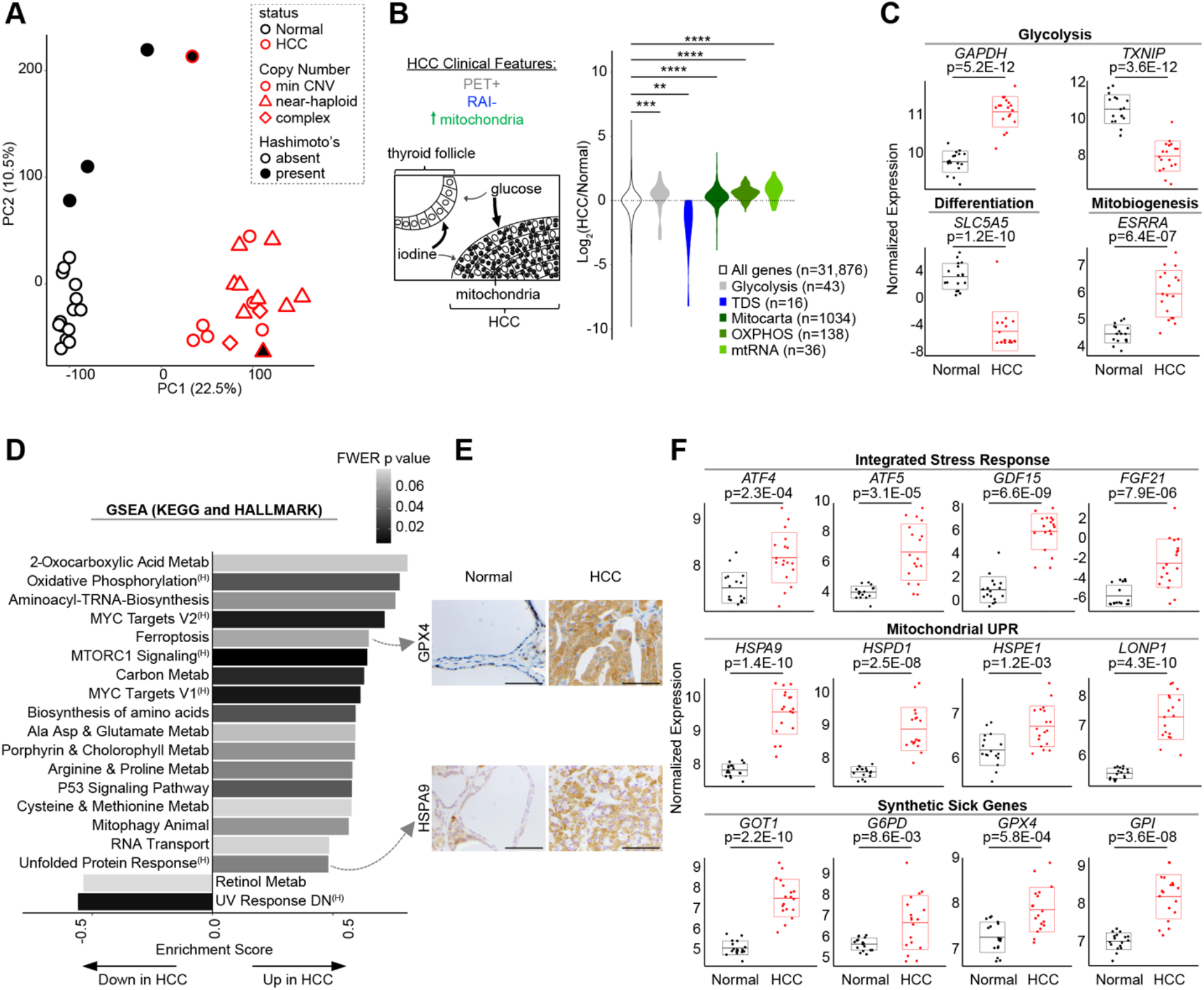
Transcriptomic landscape of HCC. (A) Principal component analysis (PCA) of HCC and normal thyroid (n=18 each) RNA-seq. Min CNV, minimum copy number variation; Hashimoto’s, Hashimoto’s thyroiditis. (B) Schematic of HCC clinical features with violin plots of gene expression fold-changes (HCC/Normal) for related molecular pathways. TDS, Thyroid Differentiation Score; mtRNA, mitochondrial RNA. Significance tested by K-S test: **p < 1.0E-03, ***p < 1.0E-04, ****p < 1.0E-05. (C) Normalized gene expression for transcripts with adjusted p values. (D) Gene Set Enrichment Analysis (GSEA) of KEGG and Hallmarks pathways ranked by enrichment score with shaded FWER values. ^(H)^, Hallmarks gene set. (E) IHC for proteins from selected pathways in HCC and normal thyroid; scale bar 200 μm. (F) Normalized gene expression for transcripts from selected pathways with adjusted p values. “Synthetic sick genes” refers to knockouts that were synthetic sick with OXPHOS dysfunction from To et al., 2019. Horizontal lines show mean, and boxes show standard deviation. See also Figure S2.

We first focused our analysis on known clinical and histologic features of HCC, including its increased ^18^fluorodeoxyglucose uptake on PET imaging, its decreased uptake of radioactive iodine, and its accumulation of mitochondria (Figure 2B) (Lowe et al., 2003; Maximo and Sobrinho-Simoes, 2000; Pryma et al., 2006). Glycolysis genes, including the redox-dependent GAPDH step, showed consistent upregulation in HCC, while TXNIP, a key negative regulator of glucose uptake with a tumor suppressor function, showed striking downregulation (Figure 2B and 2C) (Morrison et al., 2014; Parikh et al., 2007). In line with the resistance of HCC to RAI, we observed clear downregulation in the 16 genes relevant to iodine metabolism that comprise the Thyroid Differentiation Score (TDS), including dramatic downregulation of the sodium-iodide symporter, *SLC5A5* (Figure 2B and 2C) (Cancer Genome Atlas Research, 2014). We found mitochondrial transcripts, defined by the MitoCarta3.0 inventory (Rath et al., 2021), to be consistently increased in HCC, including OXPHOS and mitochondrial RNA (mtRNA) genes (Figure 2B). Furthermore, we confirmed significant upregulation of *ESRRA* (Figure 2C), a key transcription factor involved in mitochondrial biogenesis (Mootha et al., 2004), as has been reported previously in this tumor (Yoo et al., 2016).

To uncover novel transcriptional pathways present in HCC, we performed Gene Set Enrichment Analysis (Mootha et al., 2003; Subramanian et al., 2005) on the HCC transcriptome (Supplementary Table 3). We focused on gene sets from the HALLMARK and KEGG databases that were significantly altered in HCC (FWER p ≤ 0.08). We identified 18 gene sets to be significantly enriched and 2 gene sets to be significantly repressed in HCC (Figure 2D). Amongst enriched pathways, we found activation of classic oncogenic signaling (mTOR and MYC) as well as numerous pathways related to amino acid metabolism. We also found stress response programs, such as ferroptosis (an iron-dependent form of regulated cell death), mitophagy, and the unfolded protein response (UPR) to be activated in HCC (Figure 2D). Closer inspection of the UPR gene set suggested preferential activation of the mitochondrial UPR (Kampinga et al., 2009; Starck et al., 2016) in HCC, which we confirmed through a survey of all heat shock protein (HSP) members (Supplementary Figure 2A). Furthermore, GSEA on MitoCarta3.0 genes identified “protein homeostasis” as the most significantly enriched mitochondrial pathway in HCC (Supplementary Figure 2B).

We sought to validate the enrichment of select programs in HCC using immunohistochemistry (IHC). Increased immunoreactivity of GPX4 (a selenoprotein that protects against ferroptosis by scavenging lipid hydroperoxides) and HSPA9 (a member of the mitochondrial UPR) in HCC cells corroborated the transcriptional enrichment for the Ferroptosis and UPR pathways (Figure 2E, top and bottom). However, IHC for phospho-S6 (Ser235/236), a marker of mTOR activation, showed more pronounced staining in stroma and blood vessels compared to HCC cells despite strong total S6 staining throughout (Supplementary Figure 2C). This staining pattern suggests that full engagement of the mTOR response may preferentially occur in the tumor microenvironment.

The enrichment of the UPR, along with pathways related to amino acid metabolism, suggested that the integrated stress response (ISR) was activated in HCC (Mick et al., 2020). ISR transcripts were significantly increased in HCC, including *ATF4* and *ATF5* (key transcription factors coordinating the ISR), as well as *GDF15* and *FGF21* (cytokines associated with the ISR; Figure 2F, top) (Harding et al., 2003; Lehtonen et al., 2016). Moreover, as the mitochondrial UPR has been associated with ISR activation, a focused look at key mitochondrial UPR members, including *HSPA9, HSPD1, HSPE1*, and *LONP1*, confirmed coordinated upregulation of this pathway in HCC (Figure 2F, middle panel) (Fiorese et al., 2016; Khan et al., 2017).

To identify pathways that promote survival of HCC in the face of complex I loss, we considered the expression of genes previously identified from a genome-wide CRISPR-Cas9 knockout screen to cause synthetic sickness with mitochondrial dysfunction (To et al., 2019). We predicted that the expression of such “predefined synthetic sick genes” would be defended or upregulated in HCC due to mitochondrial dysfunction caused by mtDNA mutations. Consistent with our prediction, nearly all of the 14 predefined synthetic sick genes present in our RNA-seq data were upregulated, and 9 of these genes were part of enriched gene sets in HCC (Supplementary Figure 2D). It was notable that genes involved in glycolysis (*GPI*), amino acid metabolism (*GOT1*), the pentose phosphate pathway (*G6PD*), and lipid peroxide detoxification (*GPX4*) were represented as potential metabolic vulnerabilities whose expression was increased (Figure 2F, bottom).

### Metabolic landscape of HCC

In parallel, we investigated the HCC metabolome using four liquid chromatography-tandem mass spectrometry (LC-MS) methods that jointly quantified 617 known metabolites (Supplementary Table 4). The metabolome of HCC was distinct from normal thyroid with 440 known metabolites found to be differentially abundant (adjusted p value < 0.01), and PCA analysis of all known metabolites showed tumors to be distinct from normals along PC1 (32.2% of the variance; Figure 3A; Supplementary Table 4). Amongst the most significantly changing metabolites in HCC, we found those related to thyroid differentiation (thyroxine and triiodothyronine) to be decreased, while those related to glutathione (γ-glutamyl-cysteine or γ-Glu-Cys) and heme (biliverdin), were increased (Figure 3B). Consistent with the strong transcriptional changes in amino acid metabolism, we observed amino acids to be enriched in upgoing HCC metabolites (Figure 3B).

**Figure 3:**
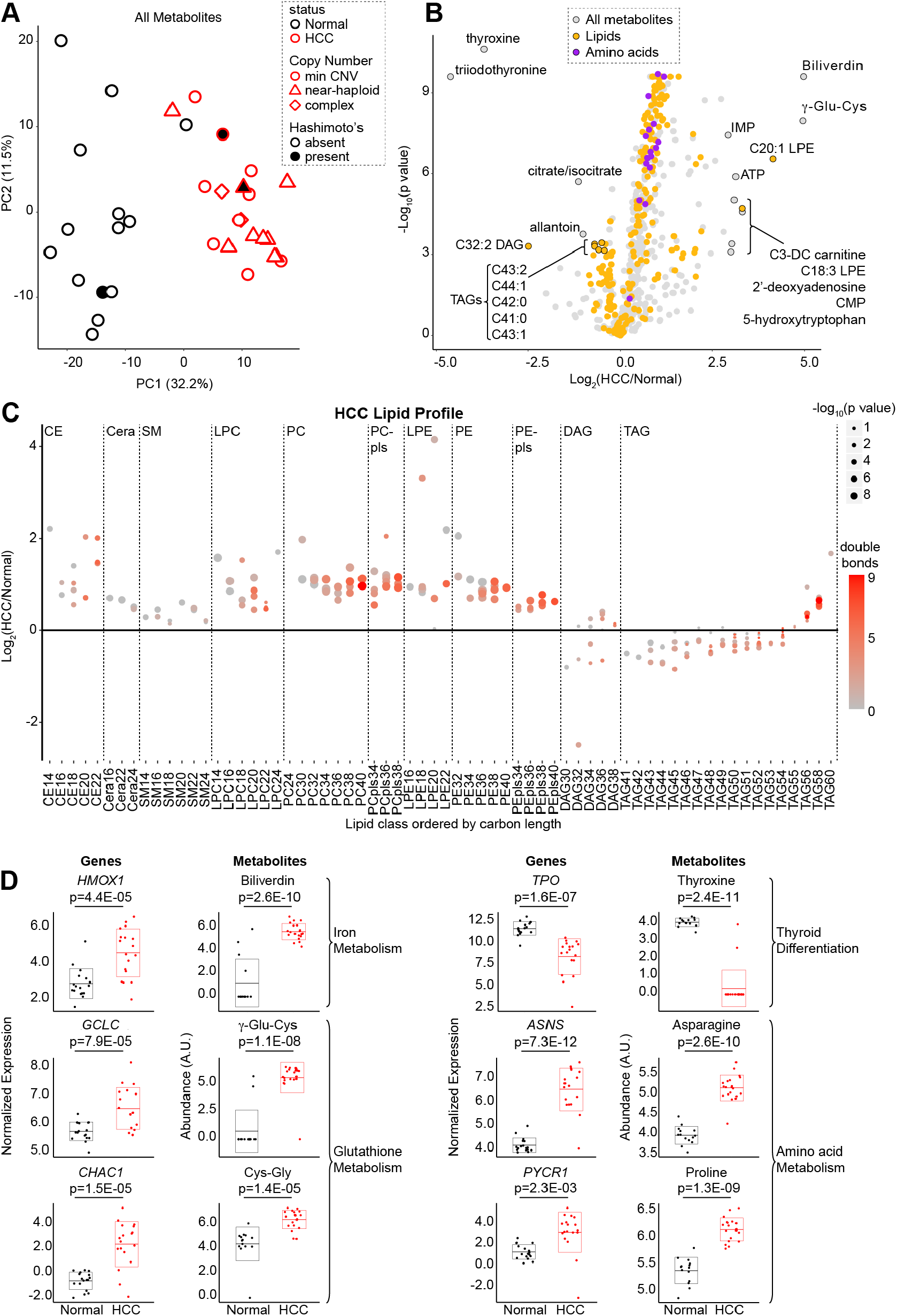
Metabolic signatures of HCC. (A) Principal component analysis (PCA) of metabolomics from four mass spectrometry methods in HCC (n=20) and normal thyroid (n=14). Min CNV, minimum copy number variation; Hashimoto’s, Hashimoto’s thyroiditis. (B) Volcano plot showing fold-change (HCC/Normal; x-axis) and adjusted p value (y-axis) with metabolites annotated by name and color. (C) Differential abundance of lipids organized by class and sorted by number of carbons with double bonds (color) and adjusted p values (dot size). CE, cholesterol ester; Cera, ceramide; SM, sphingomyelin; PC, phosphatidylcholine; LPC, lysoPC; PE, phosphatidylethanolamine; LPE, lysoPE; pls, plasmalogen; DAG, diacylglycerol; TAG, triacylglycerol. (D) Normalized expression and abundance for enzymes and their metabolic products respectively with adjusted p values. Horizontal lines show mean, and boxes show standard deviation. See also Figure S3.

A more detailed analysis of lipids in HCC revealed differential abundance of specific categories of lipid species, including an accumulation of PUFAs. In contrast to multiple triacylglycerols (TAGs) which were decreased, lysophosphatidylethanolamine (LPE) species were increased in HCC (Figure 3B). Closer inspection of lipid classes showed preferential increases in ether and glycerophospholipids as well as sterol and prenol species (Supplementary Figure 3A). When the lipids were plotted as a function of carbon and double bond number, we found similar increases in saturated, monounsaturated, and polyunsaturated levels of cholesterol ester (CE), and glycerophospholipid species (Figure 3C; Supplementary Figure 3B). On the other hand, we discovered a preferential increase of TAGs with more total carbons in HCC (Figure 3C). The elevated levels of phosphatidylethanolamines with PUFA side chains along with increased adrenate and arachidonate suggests that lipid substrates capable of ferroptosis accumulate in HCC (Figure 3C; Supplementary Figure 3C) (Kagan et al., 2017; Zou et al., 2019).

### Concordance of transcriptional and metabolomic profiles

The most differentially abundant metabolites in HCC directly connected to key transcriptional changes revealed in our RNA-seq data. The two most decreased metabolites, thyroxine and triiodothyronine, corroborate the transcriptional evidence for a non-thyrocyte state with decreased thyroid hormone production in HCC (Figure 3D). The top two increased metabolites, biliverdin and γ-Glu-Cys, are products of two transcriptionally upregulated reactions, *HMOX1* and *GCLC* respectively, that were leading edge genes in the Ferroptosis gene set (Figure 3D). We also found significant upregulation of the γ-glutamylcyclotransferase *CHAC1*, a ferroptosis biomarker, as well as its product cysteinyl-glycine (Cys-Gly), suggesting a potential state of cysteine stress in HCC (Figure 3C) (Dixon et al., 2014). A closer look at glutathione metabolism revealed increased levels of reduced and oxidized glutathione, homocysteine, and cystathionine along with transcriptional changes that suggested activation of the γ-glutamyl-cycle and stalling of transsulfuration in HCC (Supplementary Figure 3D and 3E). Finally, the three most significantly increased amino acids were alanine, asparagine, and proline, which are products of reactions identified by our GSEA analysis to be significantly upregulated, including the *ASNS* and *PYCR1* genes (Figure 3D). Collectively, our joint transcriptomic and metabolomic profiling highlights changes in glutathione, amino acid, and lipid metabolism in HCC.

### Establishing authentic HCC models

Given that the field of HCC biology has been hampered by a lack of experimental models, we sought to create patient-derived cell lines to overcome this barrier. We successfully generated a patient-derived HCC cell line using tailored culture conditions from patient HC024 (hereafter referred to as “MGH-HCC1”). We also obtained a separate HCC patient-derived xenograft (PDX) sample from the NCI’s patient-derived models repository from which we generated mouse xenografts and an additional cell line (https://pdmr.cancer.gov; distribution lot name 248138-237-R; hereafter referred to as “NCI-HCC”).

To credential MGH-HCC1 and NCI-HCC cells as authentic cellular models of HCC, we compared the mtDNA and chromosomal copy number status of each cell line to an immortalized normal thyroid cell line (Nthy-ori 3-1). Sequencing of the mtDNA identified the following: 1) a disruptive complex I mutation in MGH-HCC1 (*MT-ND1*: m.G3745A, 60.2% heteroplasmy; the same mutation identified by our original sequencing of the patient’s tumor), 2) disruptive homoplasmic mutations in two complex I genes in NCI-HCC (*MT-ND1*: m.G3916A, 99.8%; *MT-ND5*: m.CA12417C, 99.9%), and 3) no alterations in Nthy-ori 3-1 cells (Figure 4A and 1B; Supplementary Table 2). Both HCC cell lines were thereby established to be complex I mutant, while Nthy-ori 3-1 cells had wild type mtDNA. A karyotype analysis of both HCC cell lines confirmed hyperdiploid karyotypes with additional copies of chromosomes reliably retained in HCC (chr 7, 12, 20; Supplementary Figure 4A) consistent with the complex copy number state. Sequencing using anchored multiplex PCR for a panel of over 100 known cancer genes found MGH-HCC1 cells to also harbor biallelic loss of *CDKN2A* and *PTEN* and NCI-HCC cells to have truncating nonsense mutations in *NF1* and *NF2*, a hotspot *TERT* promoter variant (C228T), and partial homozygous losses of *CDKN2A* and *PTEN* (Supplementary Figure 4B).

**Figure 4:**
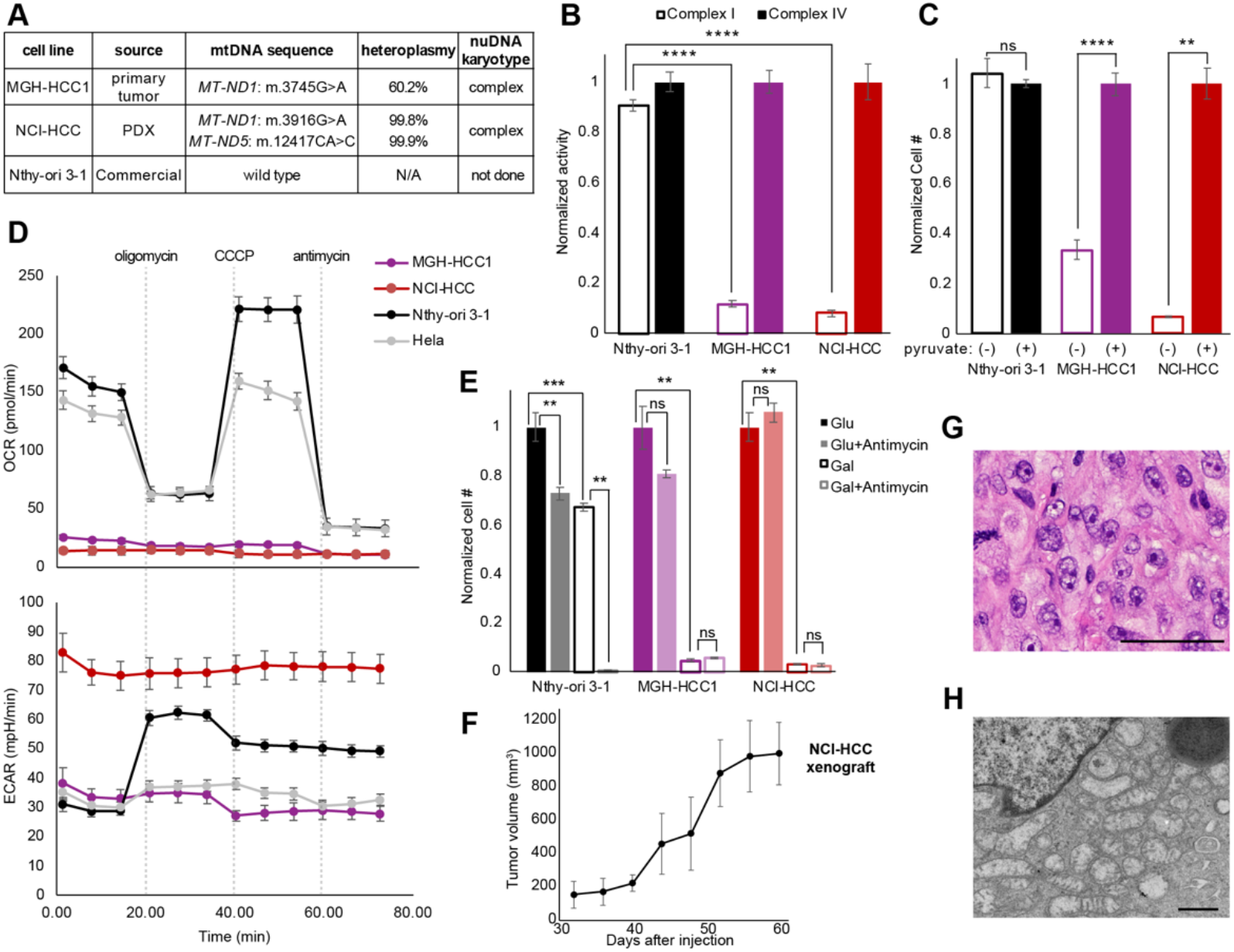
Genetics and biochemistry of authentic HCC models. (A) Table of mtDNA mutation status and nuclear karyotype of HCC (MGH-HCC1 and NCI-HCC) and normal thyroid (Nthy-ori 3-1; immortalized via SV40 transfection) cell lines. (B) Normalized enzymatic activity of complex I and complex IV from cell lines. Mean ± SD; n=3 from one experiment. (C) Normalized cell number in media ± pyruvate. Mean ± SD; n=2, representative of two experiments. (D) Oxygen consumption (OCR) and extracellular acidification rate (ECAR) of cell lines from Seahorse XFe96 analyzer. Dotted vertical lines show drug injections. Mean ± SD; n=8-10, representative of three experiments. (E) Normalized cell number of cell lines in glucose or galactose ± antimycin. Mean ± SD; n=3, representative of three experiments. (F) Tumor volumes (mm^3^) of NCI-HCC xenografts over time. Mean ± SD; n=3 from one experiment (G) H&E (scale bar 100 μm), and (H) electron microscopy (scale bar 800 nm) from NCI-HCC xenograft. For (B,C, and E) significance tested with unpaired t test: *p < 0.05, **p < 0.01, ***p < 0.001, and ****p < 0.0001. See also Figure S4.

Having established that both HCC cell lines harbor disruptive complex I mtDNA mutations, we next characterized their bioenergetic profiles. The activity of mitochondrial complex I, but not that of complex IV, was dramatically reduced in HCC cells, while Nthy-ori 3-1 cells showed intact activity for both complexes (Figure 4B). Consistent with the loss of complex I activity impairing mitochondrial NADH recycling into NAD^+^, the absence of pyruvate triggered a striking impairment in cell growth only in HCC cell lines (Figure 4C) (Titov et al., 2016). Since pyruvate enables cytosolic recycling of NADH into NAD^+^, the observed growth impairment in HCC cells can be attributed to insufficient NAD^+^ levels. HCC cells also exhibited an extremely low baseline and uncoupled oxygen consumption rate (OCR) in comparison to Nthy-ori 3-1 and Hela cells (Figure 4D). This defective mitochondrial respiration was associated with very high extracellular acidification rates (ECAR), suggesting increased rates of glycolysis. We next tested if HCC cells were capable of utilizing their mitochondria to generate ATP by culturing them in glucose- or galactose-containing media, since the use of galactose as the sole sugar source forces cells to rely on OXPHOS (Robinson et al., 1992). Growing the HCC cell lines in galactose confirmed their defective OXPHOS as this led to dramatic cell death comparable to that seen with antimycin (complex III inhibitor) treatment of Nthy-ori 3-1 cells in galactose (Figure 4E; Supplementary Figure 4C). The loss of OXPHOS in the when these cells were grown as the sole sugar source, Finally, NCI-HCC, but not MGH-HCC1, cells were able to form subcutaneous xenografts in NOD-SCID-IL2Rγ^-/-^ (NSG) mice (Figure 4F) with hematoxylin and eosin (H&E) staining and electron microscopy respectively showing large nuclei with prominent nucleoli and abundant dysmorphic mitochondria consistent with oncocytic pathology (Figure 4G, and 4H). Considering the scarcity of authentic HCC models, our HCC cell lines and PDX model are valuable resources for the field as they harbor the expected genetic, biochemical, and histological features of disease.

### CRISPR screen identifies selective essentialities in HCC

To prioritize pathways of greatest essentiality in HCC, we performed a targeted CRISPR knockout growth screen and directly compared the essentiality of key genes in NCI-HCC versus Nthy-ori 3-1 cells. We custom designed a CRISPR library to target 191 genes (Supplementary Table 5) nominated by our multidimensional profiling (75 genes), prior knowledge of mitochondrial disorders (95 genes), and experience with oncocytic tumors (21 genes). We also targeted 50 control genes that were not differentially expressed in our HCC RNA-seq data. For positive and negative controls, we targeted 10 essential genes as well as 150 cutting and non-cutting controls (Supplementary Table 5). Following infection and selection of the library in NCI-HCC and Nthy-ori 3-1 cells, serial passaging with sampling of cells on Day 5, 8, 11, 16, and 20 was performed (Figure 5A). The sgRNA cassette integrated in the genomic DNA was amplified from cells harvested at each time point and sequenced. The representation of each sgRNA was compared to its initial representation in cells on Day 5 to derive the relative fitness of a knockout in each cell line as previously described (To et al., 2019). The top scoring knockouts with reduced fitness in NCI-HCC cells 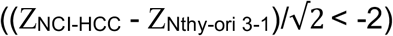 on Day 11 were *GPX4, GAPDH, GABPA*, and *G6PD*, and on Day 16 were *GPX4, GAPDH, HSPD1*, and *G6PD* (Figure 5B and Supplementary Figure 5A). Exploration of the Broad Institute’s Cancer Dependency Map showed a significant fitness reduction following *GPX4* loss in thyroid cancer cell lines, and that the thyroid lineage was the second most sensitive to *GPX4* loss of all cancer lineages tested (Figure 5C and Supplementary Figure 5B). This suggested that enhanced ferroptosis sensitivity was a general feature of thyroid cell lines, a finding supported by the decreased fitness of *GPX4* knockouts in Nthy-ori 3-1 cells albeit to a lesser extent than NCI-HCC cells. The striking dropout of *GPX4* knockouts in NCI-HCC cells reinforced that lipid peroxide stress is a vulnerability in HCC, which is consistent with our transcriptomic and metabolomic profiling.

**Figure 5:**
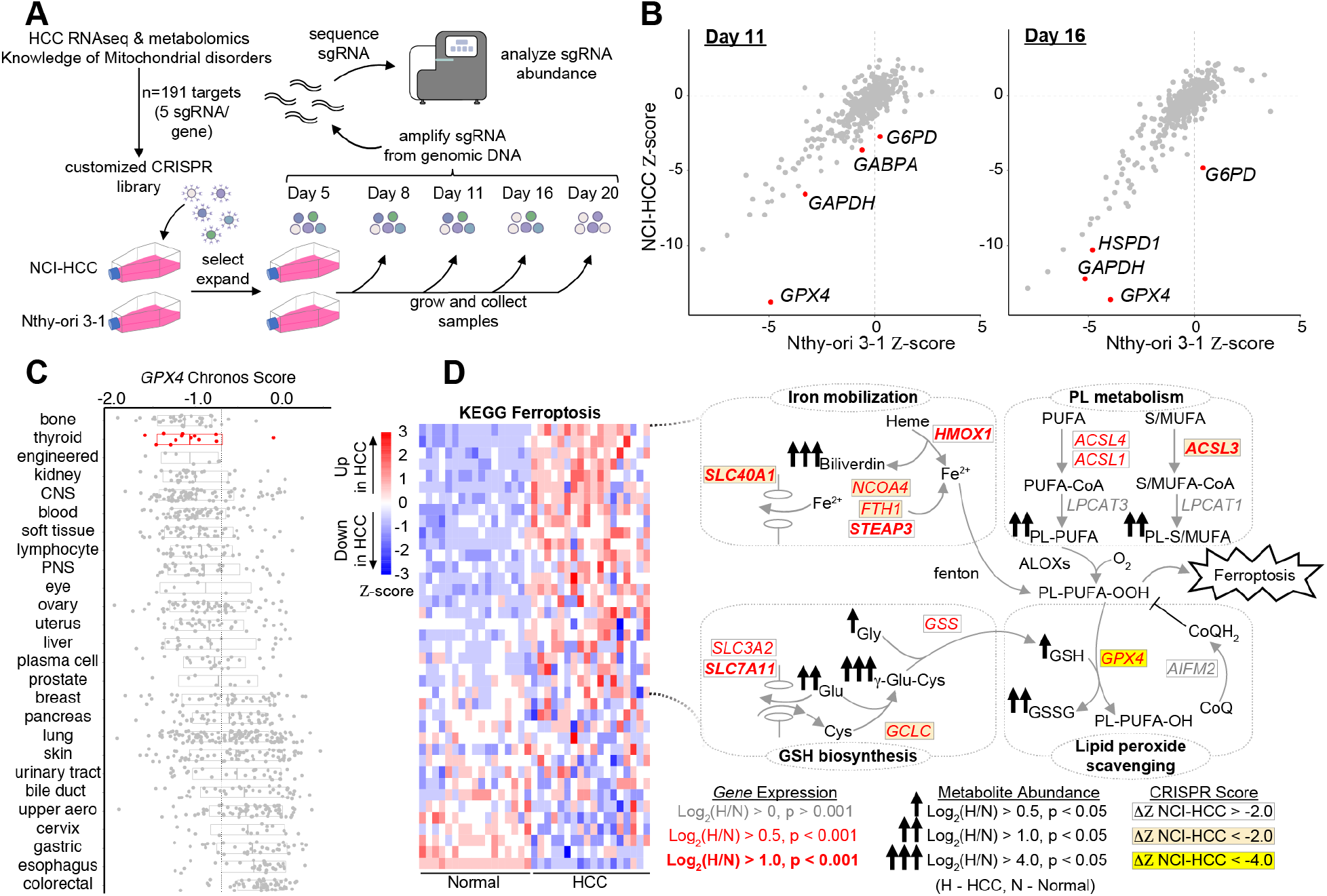
CRISPR screen identifies vulnerability to GPX4 loss in HCC. (A) Schema for customized CRISPR screen with sampling times and DNA sequencing. (B) Gene fitness scatter plots showing Z-scores in NCI-HCC (y-axis) vs Nthy-ori 3-1(x-axis) cells from Day 11 and Day 16 time points. (C) *GPX4* chronos scores from Cancer Dependency Map (DepMap) for all cancer cell lines ranked by sensitivity with mean (horizontal line). (D) Ferroptosis heatmap with schema of metabolic pathways represented in leading edge genes with fold-changes and adjusted p values in genes and metabolites as well as CRISPR fitness score annotated according to the legend. See also Figure S5.

The dependency on *GPX4* was noteworthy since the “Ferroptosis” gene set had one of the highest enrichment scores (Figure 2D), and GPX4 showed strong immunoreactivity in HCC (Figure 2E). Closer inspection of the Ferroptosis leading edge genes revealed coordinate increases in metabolic pathways relevant to the initiation, execution, and regulation of ferroptosis (Figure 5D and Supplementary Figure 5C). Genes related to phospholipid (ACSLs and *LPCAT3*) and GSH (*GCLC* and *GSS*) biosynthesis were increased concurrently with genes involved in free iron mobilization (*HMOX1, NCOA4, STEAP3, FTH1*, and *SLC40A1*) and lipid peroxide detoxification (*GPX4*; Figure 5D). An absence of a significant change in the alternate ferroptosis suppressor *AIFM2* (also known as FSP1), suggests that the GPX4 defense system may be preferentially active in HCC (Figure 5D). The observed gene expression changes were congruent with our metabolomics showing increased biliverdin (heme metabolism) and γ-Glu-Cys (GSH metabolism) together with elevated PUFA-containing phospholipids (Figure 5D). The equivalent upregulation of *ACSL4*, a gene required for ferroptosis, and *ACSL3*, a gene that protects against ferroptosis, suggests balanced incorporation of PUFAs as well as saturated and monounsaturated fatty acids (MUFAs) into membrane phospholipids (Figure 5D) (Doll et al., 2017; Soula et al., 2020).

Furthermore, the significant increase in the expression of system x^c-^ members (*SLC7A11* and *SLC3A2*) underscores the importance of cystine import for GSH biosynthesis, especially since intracellular cysteine production via transsulfuration was not activated in HCC (Figure 5D and Supplementary Figure 2E).

### HCC cells are vulnerable to GPX4 inhibition

We validated the sensitivity of HCC cells to *GPX4* loss using drugs that impair glutathione biosynthesis *in vivo* (sulfasalazine) or directly inhibit GPX4 *in vitro* (1S,3R-RSL 3, hereafter referred to as RSL3) (Figure 6A). Dose response curves testing the growth impact of RSL3 in MGH-HCC1, NCI-HCC, Nthy-ori 3-1, and Hela cells showed increased sensitivity in the three thyroid cell lines relative to Hela cells (Figure 6B). Moreover, MGH-HCC1 and NCI-HCC cells showed a significantly enhanced response to RSL3 compared to Nthy-ori 3-1 cells (Figure 6B). The toxicity induced by RSL3 could be completely rescued by the ferroptosis inhibitor, ferrostatin-1, suggesting an on-target effect (Figure 6C). In the absence of a GPX4 inhibitor with good *in vivo* bioavailability, we turned to sulfasalazine, a system x^c-^ inhibitor, to establish the feasibility for targeting lipid peroxide metabolism in our NCI-HCC PDX model. NCI-HCC xenografts were treated with either vehicle or sulfasalazine (100 mg/kg) by intraperitoneal (IP) injection once daily after reaching ∼100 mm^3^ in size. Although the effect on tumor growth was moderate, it was notable that sulfasalazine, even as a single agent, significantly decreased the pace of tumor growth with no effect on mouse body weight (Figure 6D). Furthermore, H&E analysis of tumors revealed that xenografts treated with sulfasalazine had increased levels of fibrosis/sclerosis, which corroborates that cysteine deprivation induced a tumor response in HCC (Figure 6E).

**Figure 6:**
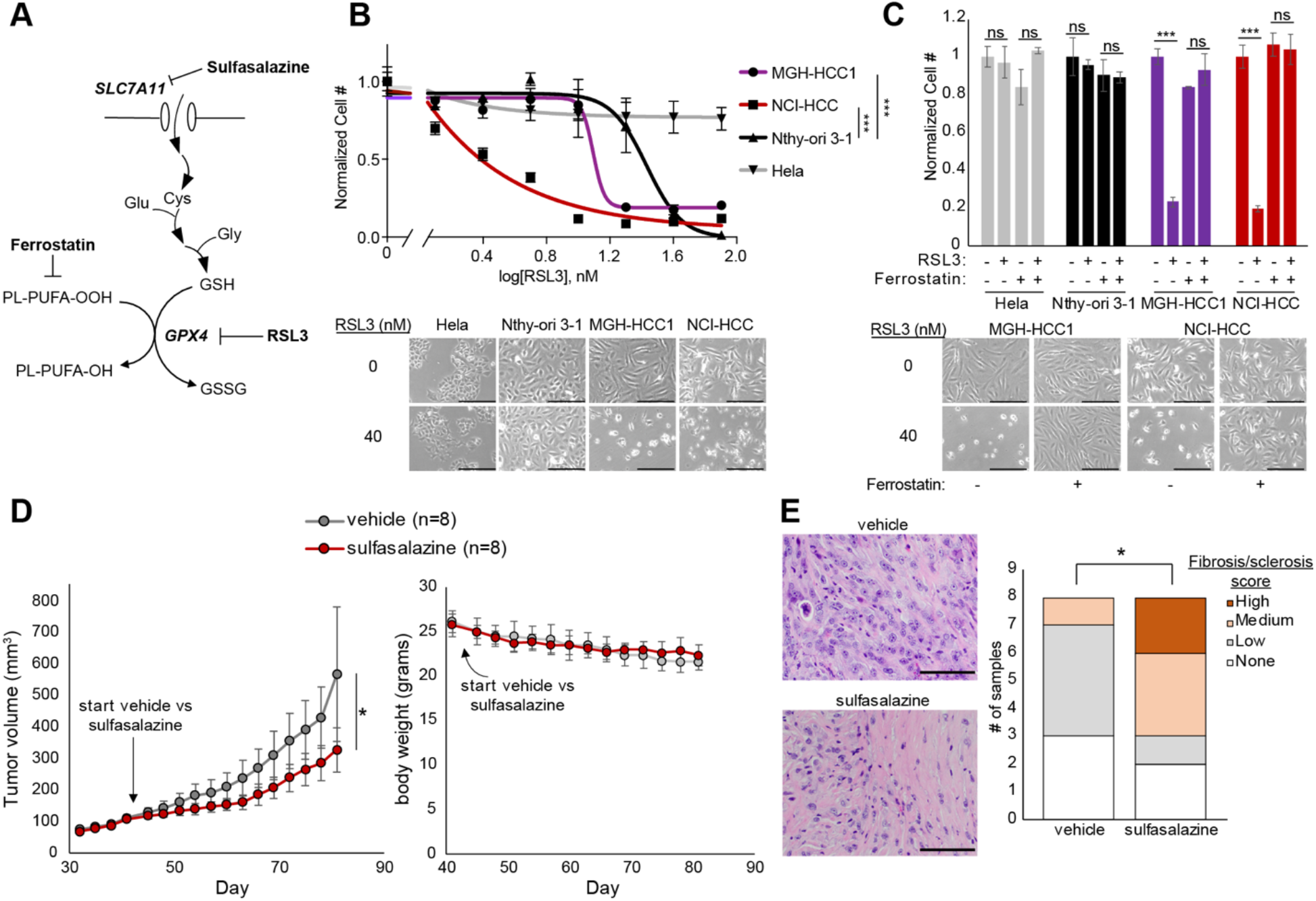
HCC cells are sensitive to ferroptosis. (A) Diagram showing glutathione (GSH) production and GPX4 reaction with drug targets. (B) Dose response curves for RSL3 plotted as normalized cell number for all cell lines with brightfield microscopy. Break in the x-axis to allow visualization of 0 nM RSL3. Scale bar 250 μm; Mean ± SD; n=3, representative of three experiments; significance tested with extra sum-of-squares F test: ***p < 0.0001. (C) Normalized cell number for cell lines with RSL3 ± ferrostatin (1 μM) with brightfield microscopy. Scale bar 250 μm; Mean ± SD; n=3, representative of two experiments; significance from unpaired t test: ***p < 0.001; ns, p > 0.05. (D) Tumor volume (mm^3^) measurements of NCI-HCC xenografts treated with vehicle (5% DMSO) or 100 mg/kg sulfasalazine via daily IP injection; Mean ± SD; n=8; significance from unpaired t test: *p < 0.05. (E) H&E of vehicle and sulfasalazine treated tumors with bar plots of fibrosis/sclerosis score. Significance tested with two-tailed test of proportions: *p < 0.05.

### Complex I loss enhances sensitivity to ferroptosis

We next sought to determine whether it was specifically the absence of complex I activity that sensitized HCC cell lines to GPX4 loss. Since HCC has two broad genetic defects, namely complex I loss and widespread LOH, the vulnerability of HCC cells to GPX4 loss could in principle be mechanistically coupled to either genetic event. Prior work has suggested that electron transport chain inhibition can modulate sensitivity to GPX4 inhibition (To et al., 2019). We utilized genetic (NDI1) and pharmacological (piericidin) perturbations to explore the relationship of complex I loss and ferroptosis (Figure 7A). For our genetic approach, we stably expressed the yeast NADH dehydrogenase, NDI1 (a single polypeptide), in the mitochondria of HCC and control cell lines to test if restoring the redox activity of complex I modulated the RSL3 response (Yagi et al., 2006). NDI1 expression in NCI-HCC cells rescued them from pyruvate auxotrophy (Figure 7B), illustrating that mitochondrial NAD^+^ production was repaired, and normalized their OCR and ECAR levels, confirming that NDI1 was functioning appropriately (Figure 7C). Consistent with complex I loss being a *bona fide* driver event in HCC, we also observed a significant reduction in cell growth in media containing pyruvate when complex I activity was restored in NCI-HCC cells relative to control cell lines (Supplementary Figure 6). In terms of RSL3 sensitivity, NDI1 expression significantly right-shifted the dose response of NCI-HCC cells, suggesting that restoration of mitochondrial complex I activity augmented resistance to GPX4 inhibition (Figure 7D). We next tested if acute inhibition of complex I activity was sufficient to confer increased sensitivity to GPX4 inhibition in a thyroid cell line with wild-type complex I. Specifically, we treated Nthy-ori 3-1 cells with piericidin, a small molecule inhibitor of complex I, in the context of RSL3 exposure. In line with complex I activity being mechanistically coupled to ferroptosis, acute inactivation of complex I with piericidin markedly sensitized Nthy-ori 3-1 cells to RSL3 (Figure 7E). This effect was directly due to complex I inhibition because NDI1 expression was sufficient to rescue the piericidin response. This rescue implicates mitochondrial NADH oxidation and downstream engagement of the respiratory chain in the regulation of ferroptosis (Figure 7E). Collectively, the ability of NDI1 to decrease the sensitivity of NCI-HCC cells to GPX4 inhibition – and the ability of piericidin to sensitize Nthy-ori 3-1 cells to GPX4 inhibition – provide gain of function and loss of function evidence that causally links complex I loss to lipid peroxide sensitivity.

**Figure 7:**
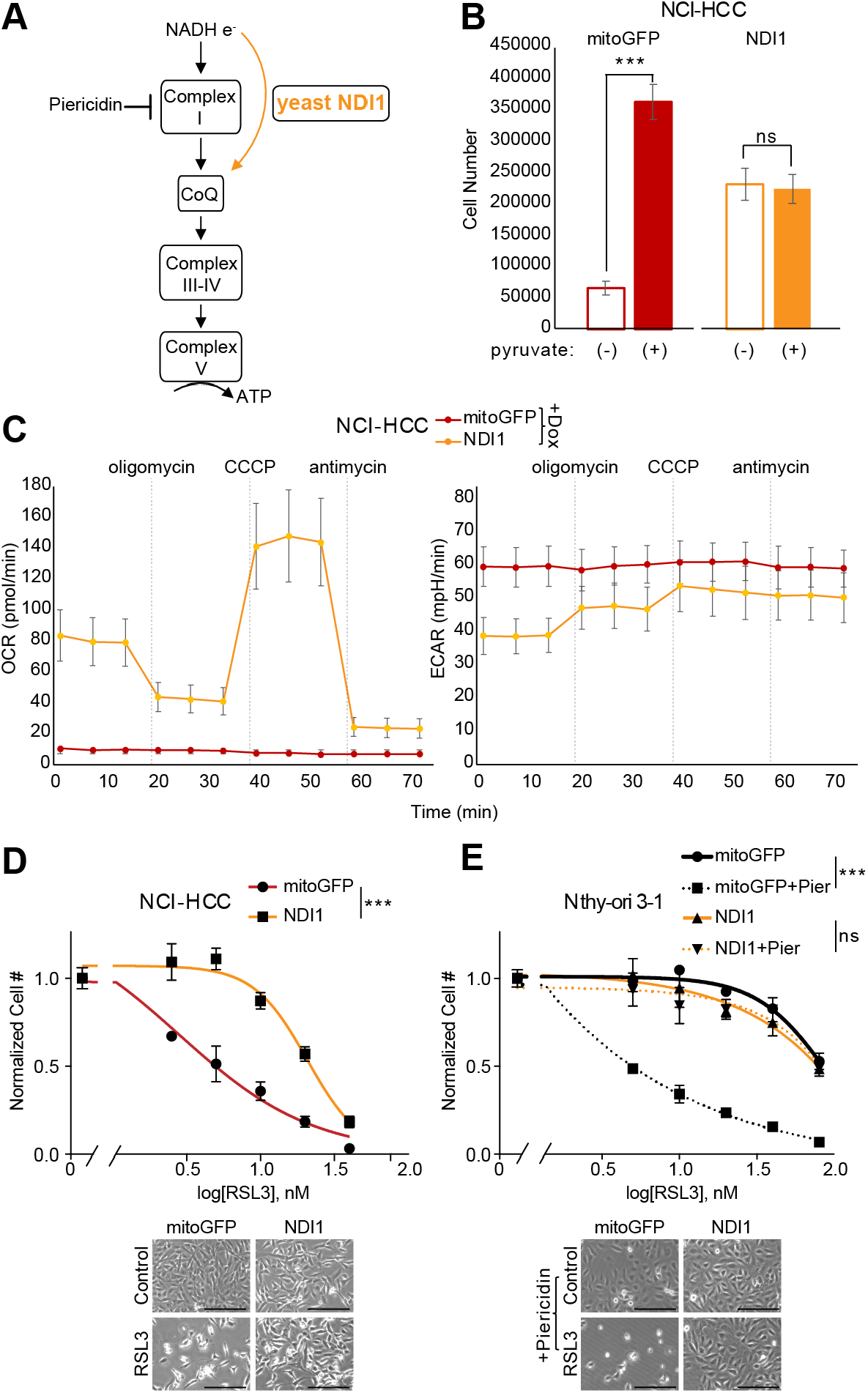
Complex I regulates sensitivity to ferroptosis. (A) Schematic of electron (e^-^) flow from NADH into OXPHOS complexes with site of action for NDI1 (yellow arrow) and piericidin (dotted line). CoQ, coenzyme Q. (B) Cell growth for NCI-HCC cells expressing mitoGFP or NDI1 ± pyruvate. Mean ± SD; n=2, representative of three experiments. Significance tested with unpaired t test: ***p < 0.001; ns, p > 0.05. (C) Seahorse XFe96 analyzer traces of oxygen consumption (OCR) and extracellular acidification rate (ECAR) in NCI-HCC cells expressing mitoGFP or NDI1 with drug injections (dotted vertical lines). Mean ± SD; n=5 representative of two experiments. (D) Dose response curves for RSL3 plotted as normalized cell number for NCI-HCC cells expressing mitoGFP or NDI1 with brightfield microscopy images. Scale bar 250 μm; Mean ± SD; n=2, representative of three experiments. (E) Dose response curves with normalized cell number for RSL3 ± piericidin in Nthy-ori 3-1 cells expressing mitoGFP or NDI1 with brightfield microscopy images. Scale bar 250 μm. Mean ± SD; n=2, representative of three experiments. For (D) and (E), break in the x-axis is to allow visualization of 0 nM RSL3, and significance tested with extra sum-of-squares F test: ***p < 0.0001. See also Figure S6.

## Discussion

Our integrated analysis of the transcriptomics and metabolomics of HCC delineates the landscape of metabolic adaptations in these tumors. The authentic HCC cellular models described in this study validate that complex I mtDNA mutations lead to severe OXPHOS dysfunction and capture the bioenergetic profile of HCC cells. Our CRISPR-Cas9 genetic screen identifies molecular effectors that are essential in HCC and discovers a vulnerability for ferroptosis due to mitochondrial complex I loss.

The transcriptional signatures of HCC relate to well-known clinical features, including enhanced PET avidity, mitochondrial accumulation, and RAI resistance. The upregulation of genes related to glycolysis (*GAPDH*) and mitochondrial biogenesis (*ESRRA*) together with the downregulation of genes involved in iodine metabolism (*SLC5A5*) provide a molecular framework for these clinical properties. The increased *GAPDH* levels are noteworthy since this glycolytic step requires NAD^+^, which likely becomes limiting in the context of “reductive stress” (high NADH/NAD^+^ ratio) following complex I loss. The activation of mitochondrial biogenesis via increased *ESRRA* expression may work in concert with defective mitophagy stemming from complex I loss to drive the iconic display of mitochondria seen in HCC(Joshi et al., 2015; Thomas et al., 2018). The enhanced mitochondrial biogenesis combined with the transcriptional features of mitochondrial dysfunction, such as increased *GDF15* and *FGF21*, herald a mitochondrial state in HCC reminiscent of the ragged-red fiber found in mitochondrial disorders (Black et al., 1975; Khan et al., 2017).

The molecular alterations revealed by our study spotlights novel programs related to growth and metabolism that are enriched in HCC. The upregulation of mTOR and MYC implicates these classic oncogenic signaling paradigms as mechanisms for growth in a tumor with mtDNA mutations and chromosomal losses. The enhanced amino acid, iron, GSH, and phospholipid metabolism in HCC likely creates a metabolic mixture that further favors growth by providing biosynthetic intermediates. The activation of stress response pathways, such as the mitochondrial UPR and lipid peroxide scavenging, could act in concert with this metabolic reprogramming to protect against complex I malfunction. As complex I defects are prone to generating ROS, stimulation of GSH biosynthesis, which is what we observed in renal oncocytoma (Gopal et al., 2018a), may boost the ability of HCC to neutralize radical species and mitigate glutamate toxicity (Arias-Mayenco et al., 2018; Kang et al., 2021).

The newly created HCC models that recapitulate defining genetic and metabolic disease features allowed us to dissect the contributions of activated molecular pathways with respect to complex I function. These models will be a valuable resource for the field moving forward, since only a single HCC cell line (XTC.UC1) with conflicting features has been reported to date (Zielke et al., 1998). Our HCC cell lines display the anticipated metabolic consequences of complex I loss with their dependence on pyruvate and glucose for growth highlighting that both reductive stress and the Warburg effect are present in HCC (Arroyo et al., 2016; King and Attardi, 1989).

Our CRISPR-Cas9 screen allowed us to determine which molecular changes are required for cell fitness in HCC. This targeted screen revealed that *GPX4* loss is a selective vulnerability in HCC, a finding that resonates with the increased iron mobilization and PUFA phospholipids identified by our -omics profiling (Yang et al., 2016). As PUFAs preferentially undergo peroxidation, the upregulation of *GPX4* and GSH biosynthesis represents logical adaptations to detoxify these dangerous species (Zou et al., 2019). The dependence of HCC on GPX4 is further illustrated by the marked sensitivity of our HCC cell lines to GPX4 inhibitors. While numerous factors, such as a high mesenchymal state, impact ferroptosis sensitivity, we discovered that a combined lineage effect together with complex I loss governs ferroptosis susceptibility in HCC (Viswanathan et al., 2017). Cells of thyroid origin are more sensitive to GPX4 loss than nearly all other lineages in the Cancer Dependency Map (Figure 5C), including renal cell cancers, which have been touted for their intrinsic ferroptosis sensitivity (Zou et al., 2019). The reasons for thyroid cells preferentially relying on GPX4 are unclear but may relate to oxidative stress due to the physiologic production of ROS during thyroid hormone production (Ohye and Sugawara, 2010).

Along with the inherent lineage sensitivity, HCC displays an extreme response to GPX4 inhibition with RSL3 due to loss of complex I redox activity (Figure 7). The ability of mitochondria to modulate the RSL3 response is illustrated by NDI1 expression restoring the redox activity of complex I in NCI-HCC cells and rendering them more resistant to GPX4 inhibition (Figure 7D). Moreover, the increased sensitivity conferred by acute pharmacological inhibition of complex I in Nthy-ori 3-1 cells further solidifies a relationship between complex I and lipid peroxidation (Figure 7E). These experiments directly implicate mitochondrial NADH oxidation and coenzyme Q (CoQ) reduction (CoQH_2_) as proximal effectors in the regulation of ferroptosis. As NDI1 expression slows the growth of NCI-HCC cells, but also renders them more resistant to ferroptosis, there appears to be a trade-off between augmented growth and susceptibility to lipid peroxide stress when complex I is lost.

The CoQ cycle acts as an additional defense system against ferroptosis that acts in parallel to GPX4. The production of CoQH_2_ at the plasma membrane and mitochondrial inner membrane, via FSP1 and DHODH respectively, has been shown to serve as a compartment-specific radical-trapping safeguard against ferroptosis (Bersuker et al., 2019; Doll et al., 2019; Mao et al., 2021). The CoQH_2_ system becomes the primary defense system when GPX4 is inhibited with the relative importance of the plasma membrane and mitochondrial CoQ pool depending on where lipid peroxides are formed. Our study complements the work of Mao et al. in establishing a role for enzymes capable of donating electrons to CoQ (DHODH and complex I) in modulating ferroptosis sensitivity. As the major flow of electrons into the mitochondrial CoQ pool is typically driven by complex I (Shaham et al., 2010), it is important to consider this first step in mitochondrial respiration as a mechanism for fine tuning the response to lipid peroxides. The exquisite sensitivity of HCC to GPX4 inhibition may result from complex I loss decreasing CoQH_2_ levels and thereby increasing the reliance on the mitochondrial GPX4 isoform to cope with the mitochondrial membrane stress associated with HCC.

Emerging data establish a clear role for GPX4 in mitochondria (Kremer et al., 2021; To et al., 2019) despite ferroptosis being considered as a distinct form of cell death independent of the mitochondrial respiratory chain (Dixon et al., 2012). In addition to the long isoform of GPX4 localizing to the mitochondrial inner membrane, GPX4 has been reliably found in the mitochondria of all 14 mouse tissues as reported in MitoCarta3.0 (Arai et al., 1996; Rath et al., 2021; To et al., 2019). Although earlier studies suggested that mitochondria are not required for ferroptosis, this notion was based on observations from cells with depleted mtDNA or activated mitophagy (Dixon et al., 2012; Gaschler et al., 2018). Subsequent data has shown that electron transport chain activity promotes ferroptosis during cysteine depletion (Gao et al., 2019), and work from our lab illustrates that electron transport chain inhibition creates a strong dependence on GPX4 (To et al., 2019). This interaction of the respiratory chain and GPX4 along with the findings of these studies implicate mitochondria in ferroptosis regulation with the extent of this contribution deriving from an interplay of genetics, lineage, and compartmental source of membrane peroxidation (Mao et al., 2021).

Our work highlights the selective importance of GPX4 for maintaining cell fitness in HCC. We directly connect the essentiality of GPX4 to the loss of mitochondrial complex I and show how modulation of the redox activity of complex I is sufficient to regulate ferroptosis sensitivity. At present, highly potent and specific GPX4 inhibitors with favorable *in vivo* PK/PD are lacking. Once developed, they could potentially be used in HCC based on our results. We also envision future studies leveraging the lineage and complex I effects that we have described in rational ways to broadly treat other types of cancers. To capitalize on the profound lineage effect that we observed, ferroptosis-inducing agents may also be an attractive strategy for other aggressive thyroid cancers. Since patients with metastatic thyroid cancer will have had all normal thyroid cells ablated, either surgically or with RAI, the lineage effect of GPX4 inhibition can be leveraged strategically without concern for toxicity to the normal thyroid gland. Finally, since complex I mtDNA mutations have been described in other cancers, genotyping the mtDNA could be an effective precision medicine strategy to nominate the use of a ferroptosis-inducer in a patient-specific manner (Gopal et al., 2018a; Grandhi et al., 2017).

## Acknowledgements

We thank the Brigham and Women’s Hospital (BWH) Cytogenomics Core laboratory, BWH Department of Pathology. CRISPR-Cas9 library was designed and produced by Adam Brown, Amy Goodale, and David Root at the Genetic Perturbation Platform at the Broad Institute. Electron microscopy was performed in the Microscopy Core of the Center for Systems Biology/Program in Membrane Biology, which is partially supported by an Inflammatory Bowel Disease Grant DK043351 and a Boston Area Diabetes and Endocrinology Research Center (BADERC) Award DK057521. We thank Ivy Rosales, Andrea Rios, and Christina Laguerre from the Mass General Histopathology Research Core Laboratory as well as Yoshiko Iwamoto from the Mass General Center for Systems Biology. This work was supported by the Bertarelli Rare Cancers Fund and a generous gift from the Elizabeth and Michael Ruane family. B.J.G. was supported by 2T32DK007028-46. S.R. was supported by NIH K00CA212468. R.K.G. was supported by NIH K12CA087723. V.K.M. is an investigator of the Howard Hughes Medical Institute.

## Author contributions

R.K.G., S.P., and V.K.M. conceived the study. R.K.G., V.R.V., S.P., and V.K.M. designed the experiments. R.K.G. and V.K.M. wrote the first draft of the manuscript with edits from all other co-authors. M.C. and R.I.S. analyzed the mtDNA. D.D.-S. analyzed the chromosomal copy number. A.P., B.R., and S.E.C. analyzed the RNA-seq and metabolomics data. V.R.V. and S.P. performed the *in vivo* experiments. S.R. and T-L.T. contributed to the CRISPR-Cas9 screen. A.S.F., M.L., Y.I., and P.M.S. performed and analyzed tissue stains. M.J.C., B.J.G., L.J.W., G.H.D. and S.P. recruited patients, acquired samples, and helped with clinical annotations. K.P. and C.C. performed the metabolomics. S.P. and V.K.M. supervised the study.

## Methods

### Resource Availability

Further information and requests for resources and reagents should be directed to and will be fulfilled by the Lead Contact, Vamsi Mootha. Cancer cell lines generated in this study are available upon request. Bulk RNA-seq and metabolomics data have been deposited at GEO and are publicly available as of the date of publication. The DOI is listed in the key resources table.

### Human Tissue Studies

HCC samples were collected and stored as part of the Mass General Brigham Institutional Review Board (protocol number 2008P001466). Frozen tissue was accessed to create the cohort used in this study and clinical annotations are included in Table S1. Frozen samples were processed for RNA-seq and metabolomics at the Broad Institute.

### In Vivo Mouse Studies

Female NSG (NOD-*scid* IL2Rγ^null^) mice of 4-6 weeks of age were purchased from Jackson Laboratory and housed at the animal care facility at Massachusetts General Hospital (MGH). All procedures were approved by the MGH Institutional Animal Care and Use Committee (IACUC) and conformed to NIH guidelines. Experimental groups were randomly assigned after tumors were established. Tumor size was tracked with digital calipers and mouse weights were recorded using a digital scale throughout the experiment.

### Cell Culture Studies

The MGH-HCC1 cell line (source: male) was derived from a resected metastasis from a patient in this cohort. The NCI-HCC cell line (source: male) was derived from a tumor grown from patient-derived xenograft material obtained from the NCI Patient-Derived Models Repository (model 248138-237-R). Nthy-ori 3-1 cell line (source: female) was purchased from Sigma and was part of the European Collection of Authenticated Cell Cultures. All cell lines were grown in humidity-controlled incubators at 21% O_2_, 5% CO_2_, and 37ºC and routinely tested for mycoplasma. MGH-HCC1 and NCI-HCC cells were routinely passaged in DMEM-11995 media with 10% fetal bovine serum (FBS) and 50 mg/ml uridine. Nthy-ori 3-1 and Hela cells (source: female) were routinely passed in RPMI-1640 media with 10% FBS. All experiments were conducted in DMEM-11995 media with 10% fetal bovine serum (FBS) and 50 mg/ml uridine unless otherwise specified.

### Mitochondrial DNA (mtDNA) sequencing

Genomic DNA (input 100 ng) isolated from HCC samples and cell lines was amplified with a two-amplicon long-range PCR method (Davis et al., 2014; Zhang et al., 2012). A TaKaRa LA Hot-Start Taq DNA polymerase was used with the following cycling parameters: 95ºC for 2 min, followed by 28 cycles of 95ºC for 20 min, 68ºC for 10 min, and concluding with 10 min at 72ºC. PCR products were purified using AMPure XP beads, sheared into 200 bp fragments with a COVARIS sonicator, and sequenced using an Illumina Miseq reagent kit v3 (150 cycles) with 75 bp paired-end reads. All reads mapping to GRCh37 chromosome MT were aligned to the revised Cambridge Reference Sequence (NC_012920) using GATK version 3.3, BWA version 0.7.12, and Picard Tools version 1.100. Somatic mtDNA variants were detected using the GATK HaplotypeCaller assuming a 20 chromosome mixture. Nonsynonymous somatic variants of potential pathological significance were identified using the following filters: 1) depth in tumor and matched normal (when available) ≥15x; 2) variant allelic fraction (VAF)>0.3 in tumor and <0.05 in matched normal; and 3) TumorVAF – Normal VAF >0.4. For tumors without matched normal tissue, we reported loss-of-function variants, the MT-TL 3244G>A variant associated with MELAS, and a missense variant in HC024 that was also reported in our whole-exome cohort as pathogenic (Gopal et al., 2018b). Variants deemed to be pathogenic were validated by RNA-seq reads for the corresponding mitochondrial RNA species using samtools mpileup and a custom R script to determine the proportions of reference and alternate alleles. A mutation was considered validated if the RNA-seq heteroplasmy was >10%. Large mtDNA deletions were identified by analyzing the uniformity of coverage for all tumor samples. For cell line mtDNA analysis, variants were considered pathogenic if they resulted in nonsynonymous changes, had a VAF>0.1, and were not associated with an mtDNA haplogroup.

### Array comparative genomic hybridization (aCGH) analysis

Genome-wide copy number variations were assessed using the SurePrint G3 Human aCGH 4×180K Microarray (Agilent Technologies, Santa Clara, CA), according to the manufacturer’s recommendations. Briefly, 1 µg of tumor DNA extracted from fresh-frozen tissue and 1 µg of male human reference DNA control (Agilent Technologies, Santa Clara, CA) were digested with *Alu*I and *Rsa*I. After restriction digestion, the sample DNA and the control DNA were individually labeled by random priming with CY5-dUTP and CY3-dUTP, respectively using the SureTag DNA labelling kit (Agilent Technologies, Santa Clara, CA). The labeled DNAs from tumors and controls were purified, mixed in equal amounts, and hybridized to the microarray for 35 to 40 hours at 65°C, using the Agilent Oligo aCGH Hybridization Kit (Santa Clara, CA). Following hybridization, the slides were washed and scanned using the Agilent G2565A Microarray Scanner (Santa Clara, CA). Microarray TIFF (.tif) images were processed with Agilent Feature Extraction Software v12.0 (Santa Clara, CA). Array data was analyzed using the Agilent CytoGenomics 2.7 software. Of note, the CytoGenomics software assumes the majority of chromosomal material in a mosaic sample is diploid. As a result, for tumor samples with a higher proportion of chromosomal haploidy than diploidy, the automatic analysis did not pass QC and yielded unusual SNP probe distributions. For these samples, we used manual reassignment of copy number values for the different aCGH Log Ratio populations and reanalyzed the data. For tumor samples with a larger proportion of haploid material, this manual approach followed by the CytoGenomics software analysis yielded the expected SNP profiles and QC metrics that fell within the preferred range.

### RNA-seq and data analysis

RNA samples were prepared and sequenced at the Broad Institute using a standard large insert strand-specific mRNA sample preparation kit (TruSeq stranded mRNA) with poly(A) selection. Total RNA was quantified using the Quant-iT− RiboGreen® RNA Assay Kit and normalized to 5 ng/μl. Following plating, 2 μL of ERCC controls (1:1000 dilution) were spiked into each sample. A 200 ng aliquot of each sample was transferred into library preparation using an automated variant of the Illumina TruSeq− Stranded mRNA Sample Preparation Kit. Following heat fragmentation and cDNA synthesis, the resultant 400 bp cDNA was then used for dual-indexed library preparation, which included ‘A’ base addition, adapter ligation using P7 adapters, and PCR enrichment using P5 adapters. After enrichment the libraries were quantified using Quant-iT PicoGreen (1:200 dilution). After normalizing samples to 5 ng/uL, the set was pooled and quantified using the KAPA Library Quantification Kit for Illumina Sequencing Platforms. Pooled libraries were normalized to 2 nM and denatured using 0.1 N NaOH prior to sequencing. Flow cell cluster amplification and sequencing were performed according to the manufacturer’s protocols using either the HiSeq 2000 or HiSeq 2500. Each run was a 101 bp paired-end with an eight-base index barcode read. RNA reads were aligned to the reference genome (hg19) using a STAR aligner and de-multiplexed and aggregated into a Picard BAM file.

After alignment, RSeQC and PicardTools BamIndexStats were run on the BAM files to assess quality of the samples. Several samples were excluded from RNA-seq analyses based on the quality control metrics. HC004_N (N, normal) had high PCR duplicates, and 5% contamination detected. Several tumor samples were excluded due to low and uneven coverage: HC001_T (T, tumor), HC005_T, HC007_T, HC010_T, HC019_T. Additionally, for transcriptomic analyses HC012_T was excluded because of the large 3.7kb deletion in the mtDNA. Raw counts were estimated from alignment files using featureCounts from the Subread package. Counts were normalized using the R package qsmooth, which normalizes assuming both global variation in samples and variation within conditions (Hicks et al., 2018). Linear regression was performed on these normalized counts after transforming with voom using the R package limma. The matrix of normalized counts was the outcome variable for the regression, and tumor/normal status was the predictor. Age, sex, and BMI were not used as covariates as qsmooth was used to quantile normalize counts and account for these differences in groups. Since all experiments were performed in a single batch, we did not correct for batch. However, we did account for inter-sample correlation introduced by the paired nature of the samples by using the individual as a blocking factor.

### GSEA

Because GSEA requires that features be comparable both within and between samples, FPKM (as calculated from raw counts) were used as input for GSEA. DESeq2 was used to transform raw counts to fpkm (Love et al., 2014). GSEA was performed for 8 databases of genesets: h.all.v7.1.symbols.gmt [Hallmarks], c1.all.v7.1.symbols.gmt [Positional], c2.all.v7.1.symbols.gmt [Curated], c2.cp.kegg.v7.1.symbols.gmt [Curated], c3.all.v7.1.symbols.gmt [Motif], c5.all.v7.1.symbols.gmt [Gene Ontology], c6.all.v7.1.symbols.gmt [Oncogenic signatures], and c7.all.v7.1.symbols.gmt [Immunologic signatures]. GSEA was also performed on a curated version of the Kegg geneset that focused on metabolic pathways: c2.cp.kegg.v7.1.symbols.gmt. Each analysis was done in the desktop version of GSEA 4.0, with 1000 permutations of phenotype labels. All other settings were as per GSEA defaults.

### Metabolomics

Fresh frozen normal thyroid and HCC samples were submitted for processing and mass spectrometry at the Broad Metabolomics platform. Tissues were weighed and homogenized in 4 volumes of water (4 µL of water/mg tissue, 4°C) using a bead beater (TissueLyser II, QIAGEN; Germantown, MD). Aqueous homogenates were profiled using four complimentary liquid chromatography tandem mass spectrometry (LC-MS) methods. Hydrophilic interaction liquid chromatography (HILIC) analyses of water-soluble metabolites in the positive ionization mode were conducted using an LC-MS system comprised of a Shimadzu Nexera X2 U-HPLC (Shimadzu Corp.; Marlborough, MA) coupled to a Q Exactive mass spectrometer (Thermo Fisher Scientific; Waltham, MA). Metabolites were extracted from homogenates (10 µL) using 90 µL of acetonitrile/methanol/formic acid (74.9:24.9:0.2 v/v/v) containing stable isotope-labeled internal standards (valine-d8, Sigma-Aldrich; St. Louis, MO; and phenylalanine-d8, Cambridge Isotope Laboratories; Andover, MA). The samples were centrifuged (10 min, 9,000 x g, 4°C), and the supernatants were injected directly onto a 150 × 2 mm, 3 µm Atlantis HILIC column (Waters; Milford, MA). The column was eluted isocratically at a flow rate of 250 µL/min with 5% mobile phase A (10 mM ammonium formate and 0.1% formic acid in water) for 0.5 minute followed by a linear gradient to 40% mobile phase B (acetonitrile with 0.1% formic acid) over 10 minutes. MS analyses were carried out using electrospray ionization in the positive ion mode using full scan analysis over 70-800 m/z at 70,000 resolution and 3 Hz data acquisition rate. Other MS settings were: sheath gas 40, sweep gas 2, spray voltage 3.5 kV, capillary temperature 350°C, S-lens RF 40, heater temperature 300°C, microscans 1, automatic gain control target 1e6, and maximum ion time 250 ms. HILIC analyses of water-soluble metabolites in the negative ionization mode were conducted using a Shimadzu Nexera X2 U-HPLC (Shimadzu Corp.; Marlborough, MA) coupled to a Q Exactive Plus mass spectrometer (Thermo Fisher Scientific; Waltham, MA). Metabolites were extracted from homogenates (30 µL) using 120 µL of 80% methanol containing inosine-15N4, thymine-d4 and glycocholate-d4 internal standards (Cambridge Isotope Laboratories; Andover, MA). The samples were centrifuged (10 min, 9,000 x g, 4°C), and the supernatants were injected directly onto a 150 × 2.0 mm Luna NH2 column (Phenomenex; Torrance, CA). The column was eluted at a flow rate of 400 µL/min with initial conditions of 10% mobile phase A (20 mM ammonium acetate and 20 mM ammonium hydroxide in water) and 90% mobile phase B (10 mM ammonium hydroxide in 75:25 v/v acetonitrile/methanol) followed by a 10 min linear gradient to 100% mobile phase A. MS analyses were carried out using electrospray ionization in the negative ion mode using full scan analysis over m/z 70-750 at 70,000 resolution and 3 Hz data acquisition rate. Additional MS settings are: ion spray voltage, -3.0 kV; capillary temperature, 350°C; probe heater temperature, 325 °C; sheath gas, 55; auxiliary gas, 10; and S-lens RF level 50.

Lipids were profiled using a Shimadzu Nexera X2 U-HPLC (Shimadzu Corp.; Marlborough, MA). Lipids were extracted from homogenates (10 µL) using 190 µL of isopropanol containing 1,2-didodecanoyl-sn-glycero-3-phosphocholine (Avanti Polar Lipids; Alabaster, AL). After centrifugation, supernatants were injected directly onto a 100 × 2.1 mm, 1.7 µm ACQUITY BEH C8 column (Waters; Milford, MA). The column was eluted isocratically with 80% mobile phase A (95:5:0.1 vol/vol/vol 10mM ammonium acetate/methanol/formic acid) for 1 minute followed by a linear gradient to 80% mobile-phase B (99.9:0.1 vol/vol methanol/formic acid) over 2 minutes, a linear gradient to 100% mobile phase B over 7 minutes, then 3 minutes at 100% mobile-phase B. MS analyses were carried out using electrospray ionization in the positive ion mode using full scan analysis over 200–1100 m/z at 70,000 resolution and 3 Hz data acquisition rate. Other MS settings were: sheath gas 50, in source CID 5 eV, sweep gas 5, spray voltage 3 kV, capillary temperature 300°C, S-lens RF 60, heater temperature 300°C, microscans 1, automatic gain control target 1e6, and maximum ion time 100 ms. Lipid identities were denoted by total acyl carbon number and total number of double bond number.

Metabolites of intermediate polarity, including free fatty acids and bile acids, were profiled using a Shimadzu Nexera X2 U-HPLC (Shimadzu Corp.; Marlborough, MA) coupled to a Q Exactive (Thermo Fisher Scientific; Waltham, MA). Homogenates (30 µL) were extracted using 90 µL of methanol containing PGE2-d4 as an internal standard (Cayman Chemical Co.; Ann Arbor, MI) and centrifuged (10 min, 9,000 x g, 4°C). The supernatants (10 µL) were injected onto a 150 × 2.1 mm ACQUITY BEH C18 column (Waters; Milford, MA). The column was eluted isocratically at a flow rate of 450 µL/min with 20% mobile phase A (0.01% formic acid in water) for 3 minutes followed by a linear gradient to 100% mobile phase B (0.01% acetic acid in acetonitril) over 12 minutes. MS analyses were carried out using electrospray ionization in the negative ion mode using full scan analysis over m/z 70-850 at 70,000 resolution and 3 Hz data acquisition rate.

Additional MS settings are: ion spray voltage, -3.5 kV; capillary temperature, 320°C; probe heater temperature, 300 °C; sheath gas, 45; auxiliary gas, 10; and S-lens RF level 60. Raw data were processed using TraceFinder 3.3 software (Thermo Fisher Scientific; Waltham, MA) and Progenesis QI (Nonlinear Dynamics; Newcastle upon Tyne, UK). For each method, metabolite identities were confirmed using authentic reference standards or reference samples.

We excluded samples from analysis for low tissue yield, or if they had poor mtDNA sequencing results (indicating general poor sample quality). Metabolites were filtered if they had less than 1000 IU average intensity across both tumors and normal samples, indicating a potentially unreliable signal. Metabolites were also filtered if they were not consistently detected in the pooled QC samples, indicating a potentially unreliable signal. Data were then log10 transformed, after addition of a very small value to avoid taking the log of 0.

The log-transformed metabolite levels were used as the outcome in a multiple multivariate regression with R package limma, using status as a predictor and age, sex, and BMI as covariates (Ritchie et al., 2015). The inter-correlation within an individual was accounted for using blocking.

### Patient-derived xenograft and cell line creation

The MGH-HCC1 cell line was derived from a resected metastasis (HC024) and cultured in DMEM-11995 media with 10% fetal bovine serum (FBS) and 50 mg/ml uridine. The tumor sample was cut into small pieces and trypsinized with pipetting to break up tissue chunks then spun at 1200 rpm for 5 min in a Beckman Coulter Allegra X-12/R centrifuge. The pellet was resuspended then passed through a cell strainer and plated into 6-well plates at 21% O_2_, 5% CO_2_, and 37ºC. Cells were then monitored with microscropy to track morphology and serially passaged with differential trysinization until an immortalized cell line with epithelial morphology resulted.

The NCI-HCC cell line derived from a patient-derived xenograft model developed by the NCI Patient-Derived Models Repository (PDMR; model 248138-237-R). A tumor grown from transplanted xenograft tissue from the NCI PDMR was harvested, cut into small pieces, and digested with collagenase. Digested material was then passed through a cell strainer and grown in DMEM-11995 media with 10% fetal bovine serum (FBS) in incubators kept at 21% O_2_, 5% CO_2_, and 37ºC. Cells were serially passaged until an immortalized cell line was established that retained the ability to form xenografts in NSG mice.

Karyotyping of the MGH-HCC1 and NCI-HCC cells lines was performed at the Brigham&Women’s Hospital CytoGenomics Core Laboratory. The karyotype was analyzed using the GTG banding pattern and counting 20 metaphase spreads with further analysis of 5 karyotypes with marker chromosome evaluation. Targeted next-generation sequencing using a customized version of the VariantPlex™ Solid Tumor Kit (ArcherDX Inc.) was performed to look for variants in 104 known cancer genes in MGH-HCC1 and NCI-HCC cells as previously described (Zheng et al., 2014). Briefly, enzymatically sheared double-stranded genomic DNA was end-repaired, adenylated, and ligated with a half-functional adapter. Next-generation sequencing using Illumina NextSeq 2×150 was performed on a sequencing library made from two hemi-nested PCR reactions.

### OXPHOS activity and oxygen consumption rate assays

Complex I and Complex IV enzyme activity assays were performed on lysates from MGH-HCC1, NCI-HCC, and Nthy-ori 3-1 cells. For each assay, 1×10^7^ cells were harvested, washed, resuspended according to the manufacturer’s (Abcam) protocol and the protein concentration was determined using BCA. Microplates were loaded with samples and incubated for 3 hours at room temperature for each assay. After washing and adding the respective assay solutions, the plates were placed in an Envision microplate reader and the enzyme activity was measured at 450 nm (complex I) and 550 nm (complex IV). Oxygen consumption rates (OCR) and extracellular acidification rates (ECAR) were measured in cell lines using a Seahorse XFe96 Analyzer (Agilent).

Cell lines were seeded at 30,000 cells/well in standard culture media. The next day, media was changed to media of the same formulation (25 mM glucose, 4 mM glutamine, and 1 mM pyruvate) except with HEPES-KOH substituted for sodium bicarbonate and with no phenol red present. Cells were placed for 1 hour into a non-CO_2_ controlled incubator at 37ºC and then transferred to the Seahorse XFe96 Analyzer for OCR and ECAR measurements. Inhibitors were added from ports to achieve the following concentrations: oligomycin 1.5 μM, CCCP 2 μM, and antimycin 0.5 μM. The same Seahorse protocol was used for NCI-HCC cells expressing mitoGFP or NDI1 except doxycycline (1 μg/mL) was added to the media.

### Cell growth and drug testing assays

For growth assays testing the effect of pyruvate, MGH-HCC1, NCI-HCC, and Nthy-ori 3-1 cells were cultured in DMEM 11965 media with 10% dialyzed FBS, and 1 mM sodium pyruvate. Cells were then seeded in triplicate at 30,000 cells/well into 6-well plates with DMEM 11965 media with 10% dialyzed FBS ± 1 mM sodium pyruvate. Cells were then harvested after 4 days and counted. The same protocol was followed with NCI-HCC cells expressing either mitoGFP or NDI1 except cells were seeded in triplicate at 100,000 cells/well in doxycycline (0.5 μg/mL) ± 1 mM sodium pyruvate and harvested 3 days later. All cell counts were done using a Z2 Coulter Particle Count and Size Analyzer (Beckman Coulter, Brea, CA) and plotted as either normalized to the average cell number with pyruvate present for each cell line, or as the total cell number.

For growth assays testing the effect of glucose vs galactose, MGH-HCC1, NCI-HCC, and Nthy-ori 3-1 cells were seeded in triplicate at 100,000 cells/well in 6-well plates in standard passaging media. The next day, cells were washed twice with PBS and then grown in DMEM 11966-025 media with either 10 mM glucose or 10 mM galactose, 10% dialyzed FBS, 1 mM sodium pyruvate, and 50 mg/ml uridine ± 1 μM antimycin. Cells were harvested after 3 days and counted using a Z2 Coulter Particle Count and Size Analyzer (Beckman Coulter, Brea, CA).

For growth assays testing the effect of 1S,3R-RSL 3 (RSL3), MGH-HCC1, NCI-HCC, Nthy-ori 3-1, and Hela cells were seeded in triplicate at 15,000 cells/well in 24-well plates in DMEM-11995 media with 10% fetal bovine serum (FBS) and 50 mg/ml uridine. The next day, cells were exposed to serial dilutions of RSL3 or DMSO. After 3 days of drug exposure, cells were harvested and counted using a Z2 Coulter Particle Count and Size Analyzer (Beckman Coulter, Brea, CA). For ferrostatin rescue experiments, cells were plated as above and then pretreated with ferrostatin (1 μM) for ∼1.5 hours before exposing to RSL3 (40 nM). Cells were exposed to RSL3 ± ferrostatin for 3 days prior to harvesting and counting. For all growth assays, brightfield microscopy images were taken using a Qimaging QICAM FAST 1394 digital camera at 4x magnification.

### CRISPR screen

A customized CRISPR library targeting 191 genes of interest, 50 control genes, and 10 essential genes with 5 sgRNAs per gene as well as 75 non-cutting and 75 cutting controls was generated at the Broad Institute. The library cloned into an “all-in-one” (pXPR_023) vector expressing the sgRNAs and SpyoCas9. Spin-infections of NCI-HCC and Nthy-ori 3-1 cells were performed in 12-well plates with 500,000 cells/well to achieve a multiplicity of infection (MOI) of 0.3-0.5. Optimal conditions for infection were empirically determined by infecting cells with different volumes of virus (0, 60, 80, 100, 120, 160 μL) with polybrene (0.8 μg/mL) in each cell line. Cells were centrifuged for 1 hr at 931xg at 30ºC. Approximately 24 hrs later, cells were harvested and transferred to 6-well plates and exposed to either vehicle or puromycin. Cells were then counted 48 hrs later to determine the virus volume that yielded a MOI of 0.3-0.5.

A screening-scale infection was then performed in NCI-HCC and Nthy-ori 3-1 cells with the empirically determined optimal virus amount in 12-well plates (500,000 cells/well). Infections were performed to reach a library representation of ∼1000 cells/guide following puromycin selection. The day after infection, all wells from each cell line were pooled and exposed to puromycin (2.0 μg/mL) for 48 hours. After selection as complete, cells were passaged in 175 cm^2^ T flasks and genomic DNA was isolated on Days 5, 8, 11, 16, and 20 post infection using a NucleoSpin kit (Takara). PCR was performed as previously described and samples were sequenced on a MiSeq sequencer (Illumina) (Doench et al., 2016; To et al., 2019). Raw sgRNA counts were normalized and transformed according to the formula: Log_2_(reads from an individual sgRNA / total reads in the sample × 10^6^. The relative fitness of a knockout was determined according to the formula 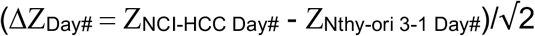, where the Z score is based on the Log_2_ fold-change of normalized sgRNA count for a given day # relative to day 5. Top scoring knockouts were identified by comparing the relative fitness in NCI-HCC and Nthy-ori 3-1 cells and defined as ΔZ_Day#_ < -2.

### Xenograft growth assays

For tumor growth assays, 2×10^6^ cells were injected subcutaneously in Matrigel into the flank of NSG mice. Mice weights were tracked with digital scales and tumor size was measured using digital calipers every 3-4 days with volumes calculated from the formula volume = ½(length x width^2^). For the sulfasalazine experiment, mice were randomized to control vs treatment groups once tumors reached ∼100 mm^3^. Vehicle (5% DMSO in saline) and Sulfasalazine (100 mg/kg in 5% DMSO in saline) were injected intraperitoneally daily. Sulfasalazine was dissolved in DMSO and diluted in saline prior to injection.

### Histology, immunohistochemistry, and electron microscopy

Formalin-fixed paraffin-embedded human normal thyroid and HCC samples as well as tumor xenografts were mounted as 5 µm sections and processed for immunohistochemistry (IHC) at the Massachusetts General Hospital (MGH) Histopathology Core. Following baking, deparaffinization, and rehydration, sections were rinsed in ddH_2_O or PBS (pH 7.4) and incubated for 10 minutes in 3% peroxide/water or 3% peroxide/methanol. Heat-induced epitope retrieval was performed in a water bath at 97°C using either 0.5M EDTA (pH 8.0), Sodium Citrate, or Borg Decloaker (BioCare Medical, BD1000, pH 9.5). Primary antibodies used were as follows: GRP75 (Cell Signaling Technology #3593, 1:50); S6 (Cell Signaling Technology #2217, 1:150, sodium citrate retrieval); Phospho-S6 ser 235/236 (Cell Signaling Technology #2211S, 1:400, sodium citrate retrieval); GPX4 (Abcam ab125066, 1:100, sodium citrate retrieval); 4-HNE (Abcam ab46545, 1:100, EDTA retrieval). Cell Signaling Technology and Abcam antibodies were diluted with SignalStain Antibody Diluent (Cell Signaling Technology, #8112) and 1% Bovine Serum Albumin/PBS, respectively. A biotinylated secondary antibody (Vector Laboratories, BA-2000) or an HRP labeled polymer was followed with a DAB substrate from Vector (Vector Laboratories, SK-4100) or with the DAKO Envision+ System (DAKO, K4000) and counterstained with Hematoxylin 2 prior to coverslipping using Permount (Fisher, SP15). H&E staining was scored in a blinded fashion. Xenografts were processed for immunofluorescence (IF) with heat induced antigen retrieval using Retrievagen A (pH 6.0, BD Biosciences) and primary antibodies as follows: Ki-67 (BD Biosciences #550609, 1:50), cleaved caspase-3 (Asp175; Cell Signaling #9661S, 1:100), and 4-HNE (Abcam ab46545, 1:100).

Biotinylated secondary antibodies (1:100, Vector Laboratories, BA-2000 or BA-1000) were followed by streptavidin DyLight 594 (1:600, Vector Laboratories) with a DAPI counterstain (1:3000) and images capture using a digital scanner NanoZoomer 2.0RS (Hamamatsu, Japan).

Electron microscopy (EM) of xenograft tissue was performed at the MGH Program in Membrane Biology microscopy core facility. Xenograft tissue pieces were fixed overnight in 2% paraformaldehyde/2.5% glutaraldehyde in 0.1 M sodium cacodylate buffer (pH 7.4), washed, infiltrated 1 hr in 1% osmium tetroxide, washed, and then dehydrated into 100% propylene oxide. Samples were incubated in a mix of propylene oxide:Eponate resin (Ted Pella, Redding, CA), then placed into a 2:1 mix of Eponate:propylene oxide (≥2 hrs), transferred into fresh 100% Eponate and polymerized in flat molds in a 60°C oven (24-48 hrs). Ultra-thin (70 nm) sections were cut using a Leica EM UC7 ultramicrotome, collected onto formvar-coated grids, stained with 2% uranyl acetate and Reynold’s lead citrate, and examined in a JEOL JEM 1011 transmission electron microscope at 80 kV. Images were collected using an AMT digital imaging system with proprietary image capture software (Advanced Microscopy Techniques, Danvers, MA).

### NDI1 expression and cell culture

A Tet-On 3G inducible expression system for mitochondrial-localized GFP (mitoGFP) and NDI1 was established in NCI-HCC and Nthy-ori 3-1 cells. The NDI1 protein contained an N-terminal FLAG tag after the mitochondrial targeting sequence (Titov et al., 2016). Both cell lines were initially infected with the pLVX-Tet3G (Clontech) regulator vector and then underwent selection with Geneticin (1 mg/mL). Surviving cells were then infected with lentivirus for either mitoGFP or NDI1 expressed in the pLVX-TRE3G (Clontech) response vector and selected with puromycin (2 μg/mL).

### Quantification and Statistical Analysis

For all analyses, significance was determined at p < 0.05. *, **, *** represent varying degrees of significance between groups as defined in figure legends. To study differences between two groups, a Student’s t test was utilized to look for significance. For dose-response curves, significant differences in the LogIC50 value were assessed using an extra sum-of-squares F test after performing a least squares regression curve fit of the data. Randomization of mouse xenografts prior to starting drug treatments was done using a randomization function in Microsoft Excel. Figure legends define n, the number of times an experiment was performed, and what center and dispersion values are shown. For *in vivo* studies, n defines the number of biological replicates and for *in vitro* studies n defines the number of replicate wells. All analyses were performed in Microsoft Excel (Version 16.54), GraphPad Prism (Version 9.3.1), or using R code. The line and box for genes and metabolites represent the mean and standard deviations.

## Supplementary Figures

**Supplementary Figure 1:**
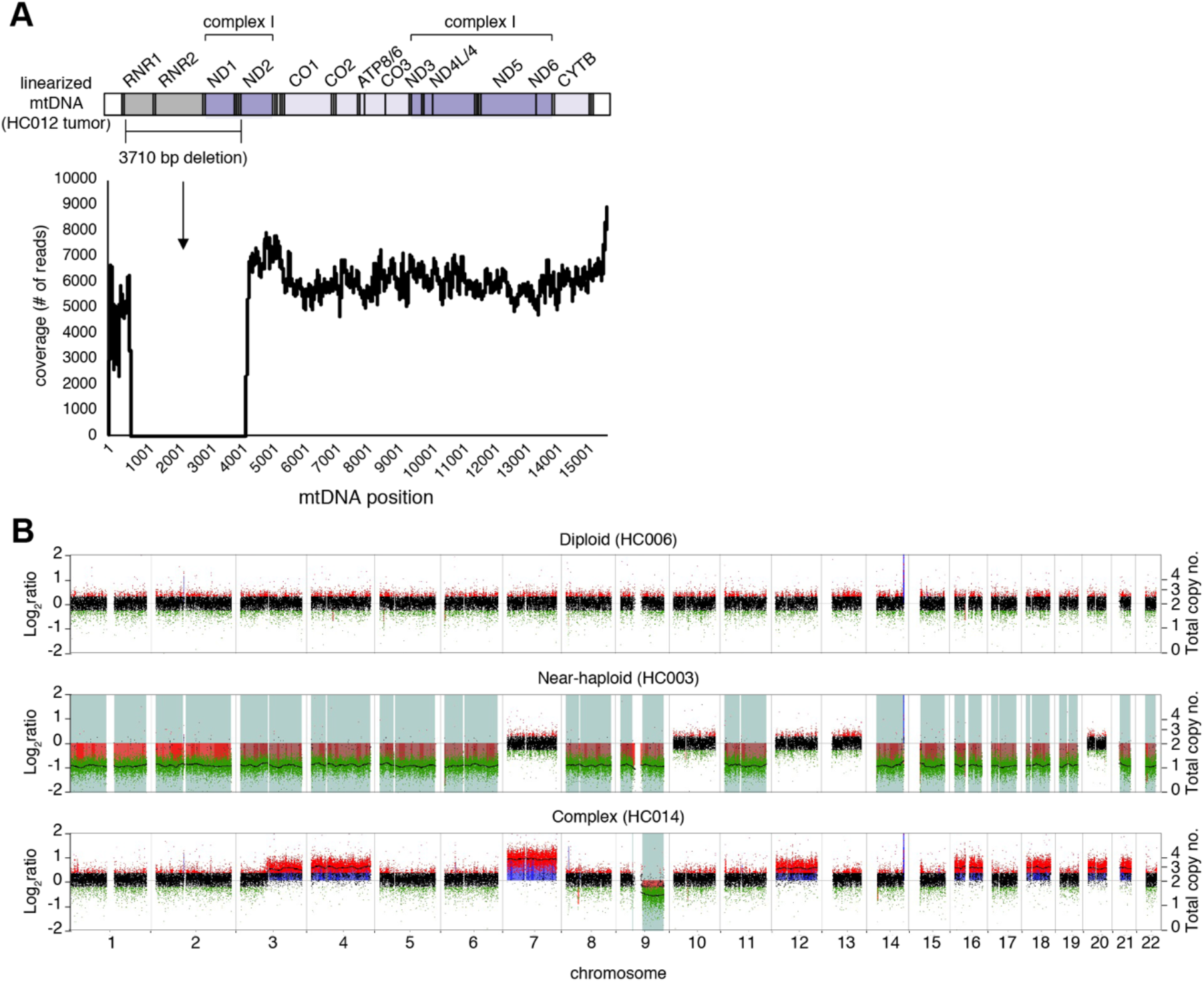
Mitochondrial and nuclear DNA alterations in an HCC cohort. Related to Figure 1. (A) mtDNA coverage in tumor sample HC012 with location of 3710 bp shown on linearized mtDNA molecule with genes labelled. (B) Genomic plots showing copy number and LOH in HCC with illustrative diploid, near-haploid, and complex copy number profiles. Log_2_ ratios (left y-axis) of the fluorescent intensities of the HCC sample vs a normal control were used to calculate copy number changes and to identify total copy number in tumor samples (right y-axis). Shaded areas indicate LOH.

**Supplementary Figure 2:**
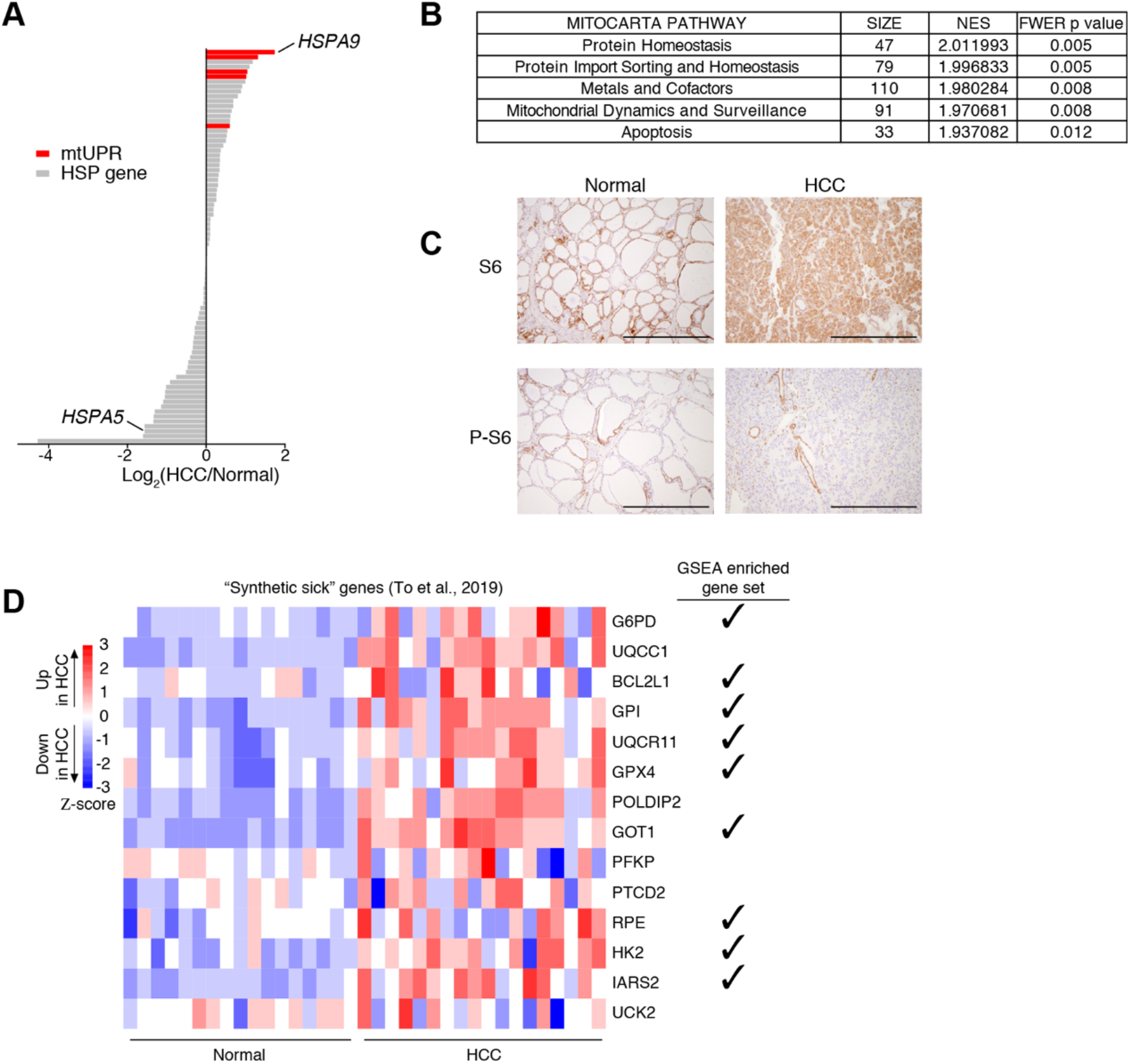
Transcriptomic landscape of HCC. Related to Figure 2. (A) Gene expression fold-change of heat shock protein (HSP) members (Kampinga et al., 2008) in HCC with *HSPA9* (a mitochondrial HSP) and *HSPA5* (an endoplasmic reticulum HSP) highlighted. (B) Gene Set Enrichment Analysis (GSEA) of MitoCarta3.0 pathways (Rath et al., 2021) with FWER p < 0.015. (C) IHC for S6 and P-S6 (phospho-S6) proteins in HCC and normal thyroid; scale bar 800 μm. (D) Heatmap of genes whose knockouts were synthetic sick with OXPHOS dysfunction (To et al., 2019) with check marks indicating members of significantly enriched KEGG&HALLMARKS gene sets.

**Supplementary Figure 3:**
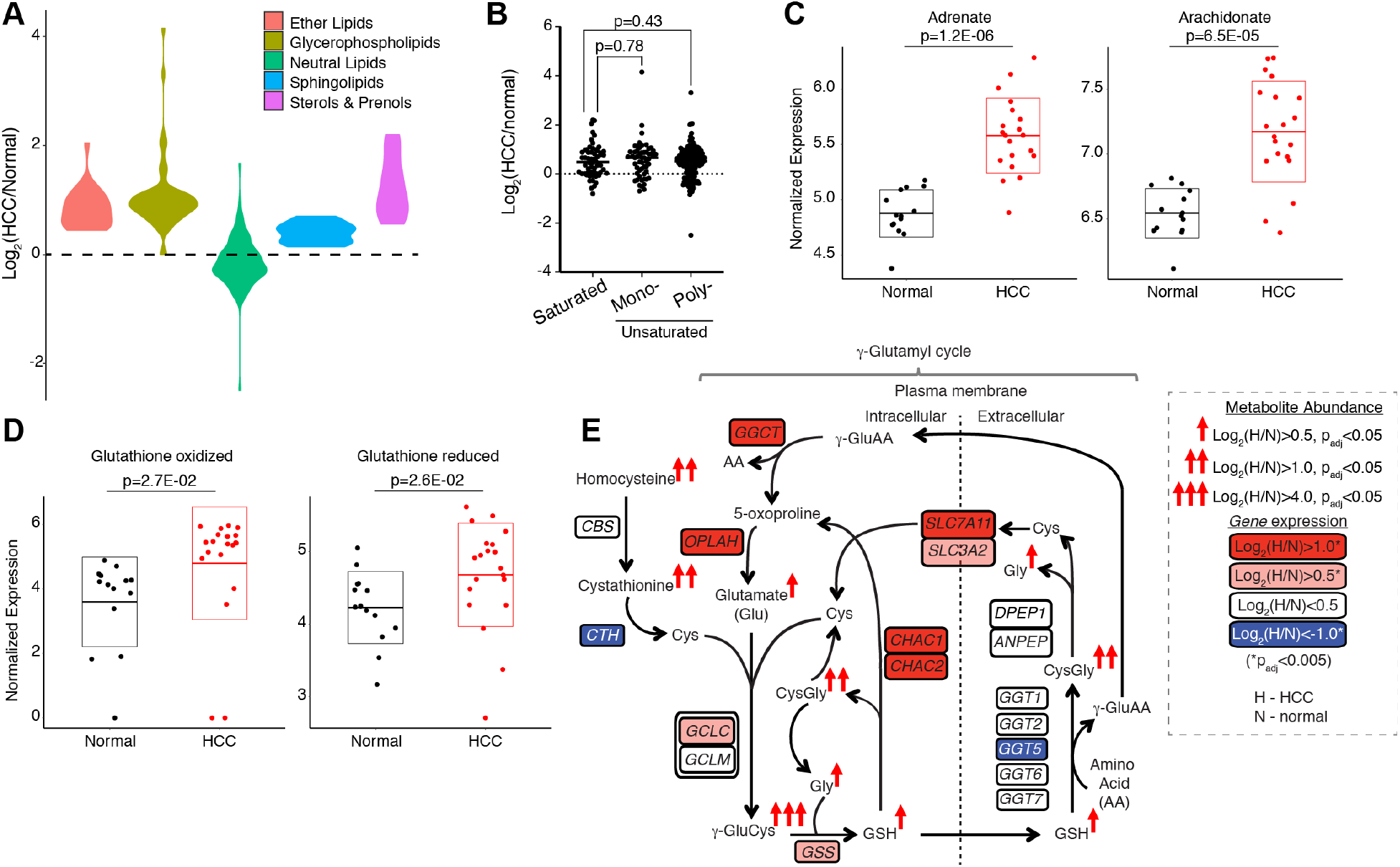
Metabolic signatures of HCC. Related to Figure 3. (A) Violin plots of gene expression fold-changes grouped by lipid category. (B) Fold-change abundance of all classes of lipids in HCC according to saturation. Horizontal bars show median; p values from two-tailed t test. (C) and (D) Normalized expression of adrenate and arachidonate as well as oxidized and reduced glutathione with adjusted p values. Horizontal bars show mean and boxes standard deviation. (E) Schema of γ-glutamyl cycle with changes in gene expression and metabolite levels according to key.

**Supplementary Figure 4:**
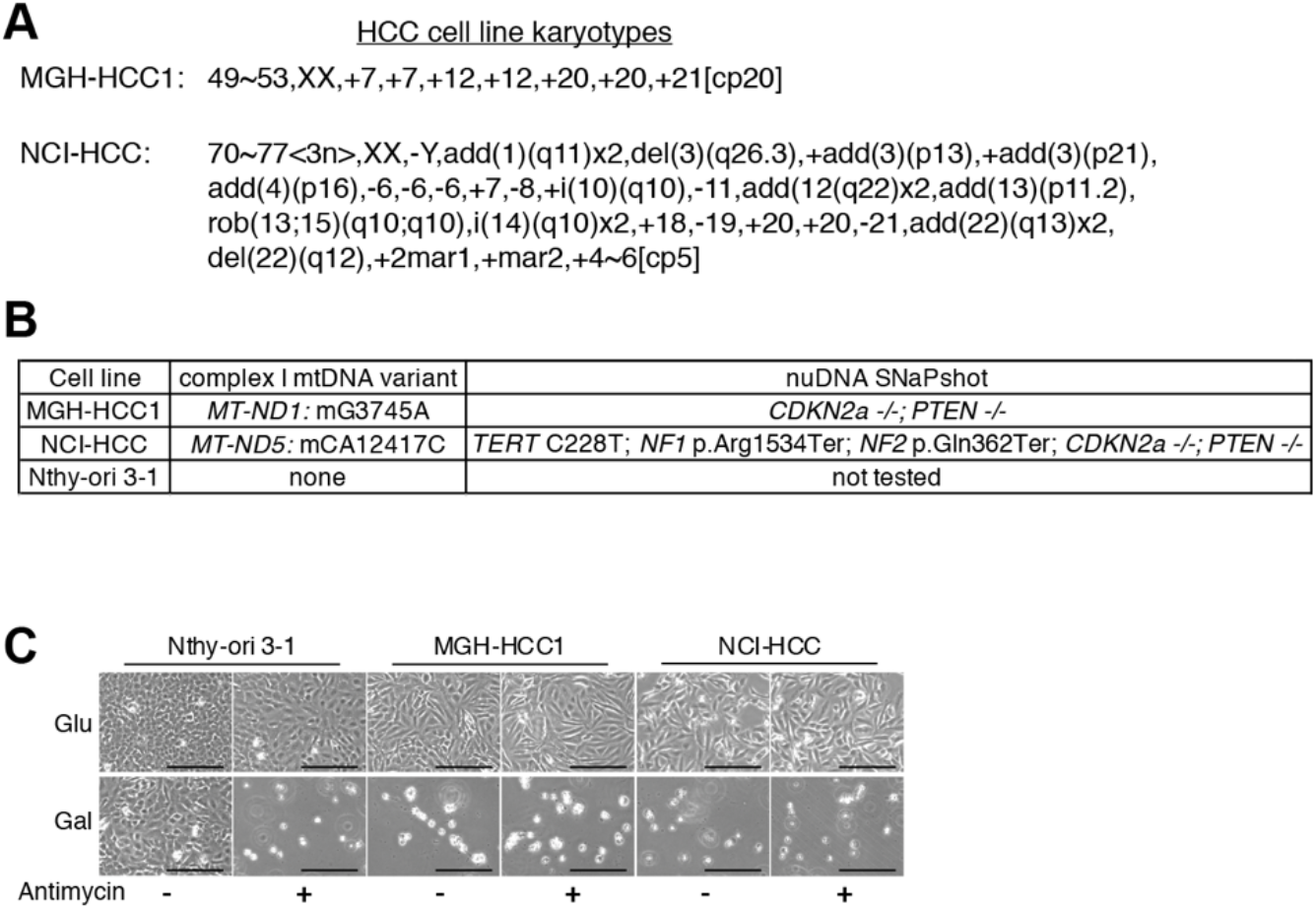
Authentic models of HCC. Related to Figure 4. (A) Cytogenomic analysis of HCC cell line karyotypes based on GTG banding of metaphase spreads. (B) Table of mtDNA and nuclear DNA (nuDNA) variants. -/-indicates biallelic loss of ≥ 1 exon. In NCI-HCC cells there was homozygous loss of exon 1 in *CDK2NA* and homozygous loss of exons 2-8 in *PTEN*, while MGH-HCC1 had complete loss of both genes. (C) Brightfield images of cell lines in glucose or galactose ± antimycine; scale bar 250 μm.

**Supplementary Figure 5:**
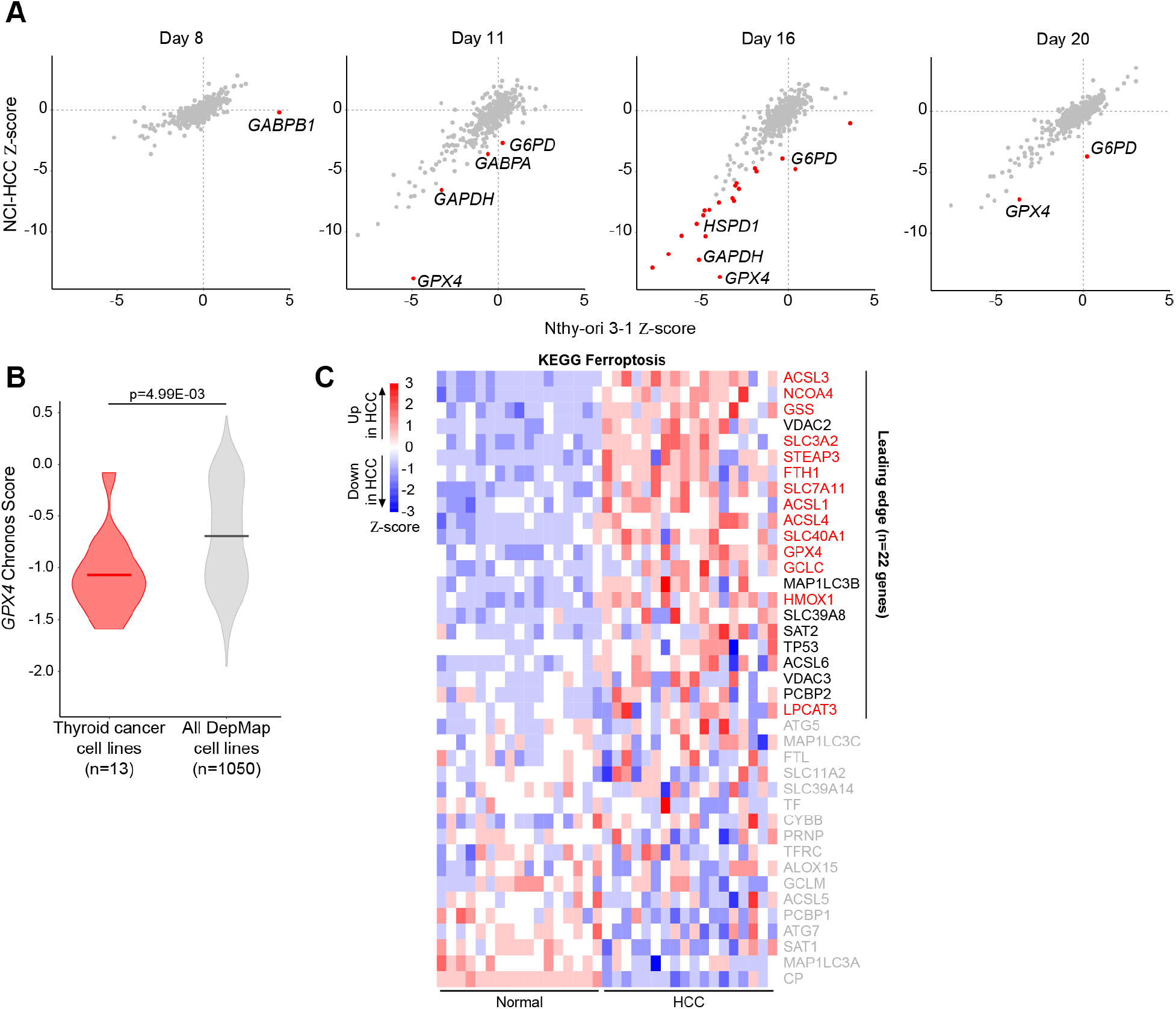
CRISPR screen identifies vulnerability to GPX4 loss in HCC. Related to Figure 5. (A) Gene fitness scatter plots showing Z-scores in NCI-HCC (y-axis) vs Nthy-ori 3-1(x-axis) cells from Days 8, 11, 16 and 20. Red dots indicate genes with ΔZ < -2 for a given day where 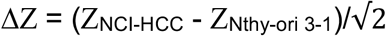 with the top 4 scoring genes labelled for Day 16. (B) Violin plots of *GPX4* chronos scores in thyroid compared to all other cancer cell lines. Horizontal lines show mean. (C) Ferroptosis heatmap with gene labels. Leading edge genes shown in red were used to create the metabolic schema in Figure 5D.

**Supplementary Figure 6:**
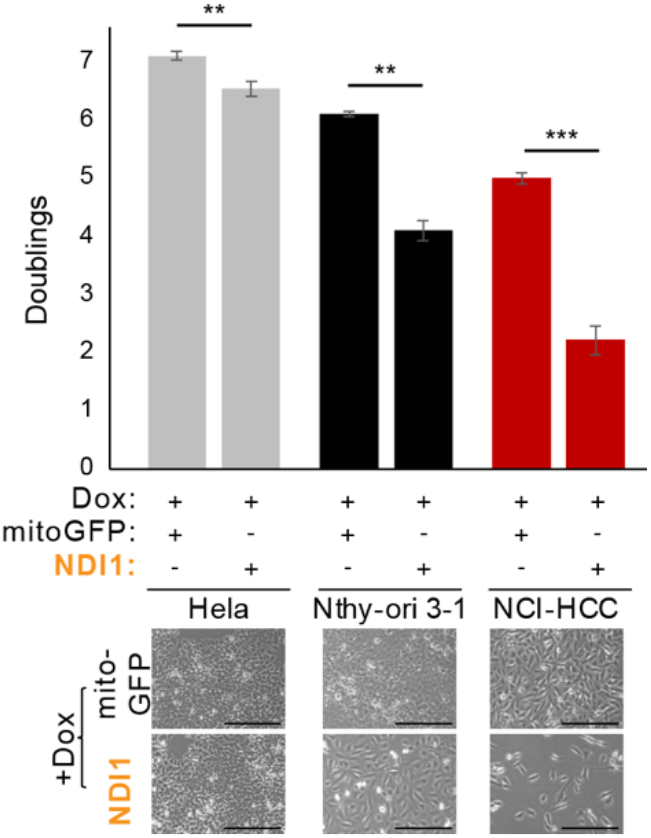
Growth effects of NDI1. Related to Figure 7. Population doublings for doxycycline-inducible mitoGFP and NDI1 cell lines with brightfield images. Mean ± SD, n=3 from one experiment; p values from unpaired t test; **p < 0.01; ***p < 0.001. Scale bar 250 μm.

